# Second messengers and divergent HD-GYP enzymes regulate 3’,3’-cGAMP signaling

**DOI:** 10.1101/600031

**Authors:** Todd A. Wright, Lucy Jiang, James J. Park, Wyatt A. Anderson, Ge Chen, Zachary F. Hallberg, Beiyan Nan, Ming C. Hammond

**Affiliations:** Department of Chemistry, University of California, Berkeley, 94720, USA; Department of Chemistry and Henry Eyring Center for Cell and Genome Science, University of Utah, Salt Lake City, UT, 84112, USA; Department of Biology, Texas A&M University, College Station, TX, 77843, USA

## Abstract

3’,3’-cyclic GMP-AMP (cGAMP) is the third cyclic dinucleotide (CDN) to be discovered in bacteria. No activators of cGAMP signaling have yet been identified, and the signaling pathways for cGAMP have appeared narrowly distributed based upon the characterized synthases, DncV and Hypr GGDEFs. Here we report that the ubiquitous second messenger cyclic AMP (cAMP) is an activator of the Hypr GGDEF enzyme GacB from *Myxococcus xanthus*. Furthermore, we show that GacB is inhibited directly by cyclic di-GMP, which provides evidence for cross-regulation between different CDN pathways. Finally, we reveal that the HD-GYP enzyme PmxA is a cGAMP-specific phosphodiesterase (GAP) that promotes resistance to osmotic stress in *M. xanthus*. A signature amino acid change in PmxA was found to reprogram substrate specificity and was applied to predict the presence of non-canonical HD-GYP phosphodiesterases in many bacterial species, including phyla previously not known to utilize cGAMP signaling.

---

Diverse signaling pathways are comprised of specific enzyme classes that serve as ‘writers’ and ‘erasers’, as well as biomolecules that serve as ‘readers’ of the signal (1). In cyclic dinucleotide signaling, ‘writer’ synthases produce the cyclic dinucleotide (CDN) signal from nucleotide triphosphate (NTP) substrates, ‘eraser’ phosphodiesterases (PDEs) degrade the CDN via hydrolysis of one or both nucleotide linkages, and ‘reader’ effectors, including transcription factors, enzymes, and riboswitches, sense the CDN signal (2, 3). So far, three CDN second messengers have been found to control critical bacterial processes, including biofilm formation, cell wall homeostasis, intestinal colonization, and transient surface interactions (4–7). Whereas representative examples of each of the three components required for signaling have been identified for both cyclic di-GMP and cyclic di-AMP, the recently discovered cyclic GMP-AMP (cGAMP) signaling pathway has remained incomplete (Fig. 1).

**Figure 1.**
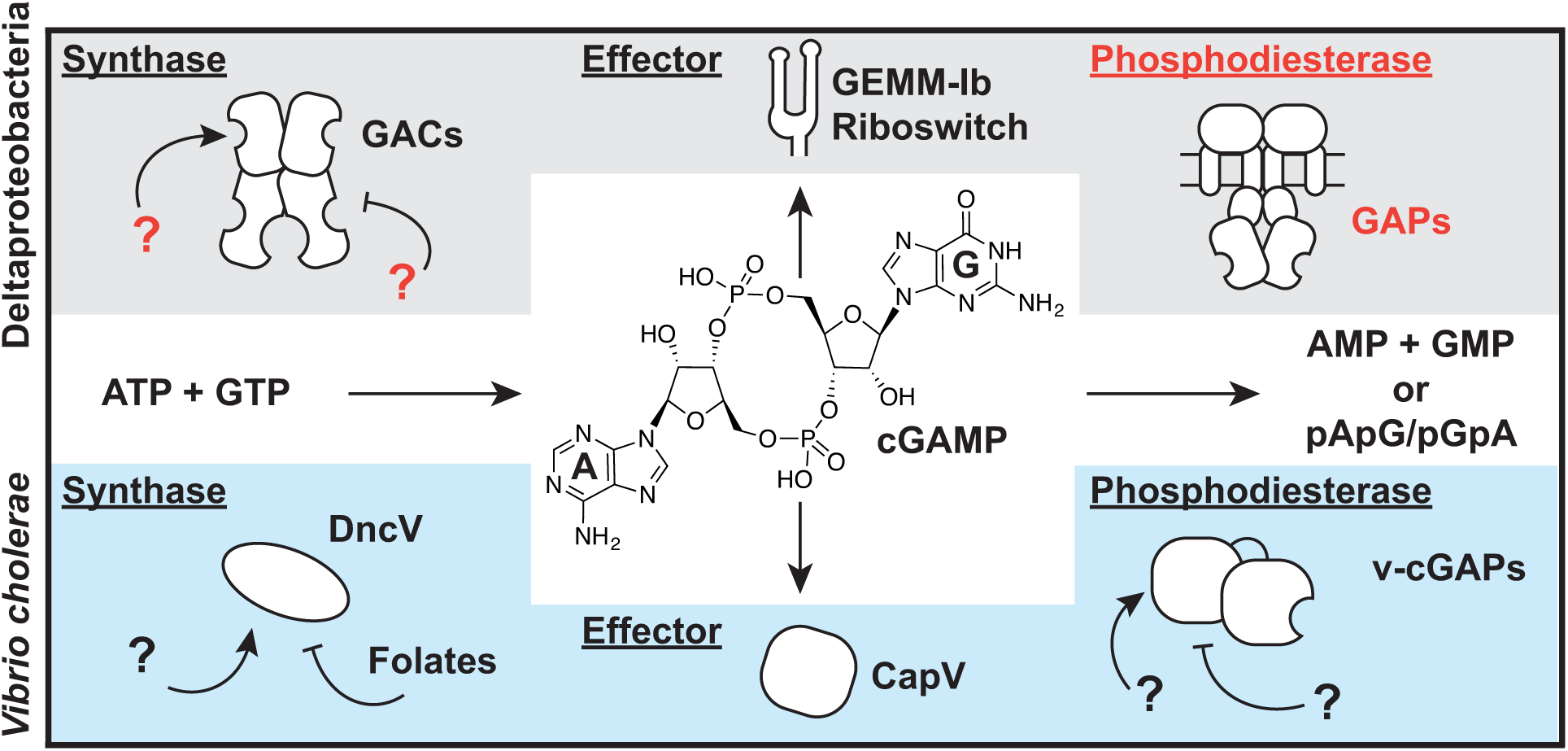
Two distinct pathways for cGAMP signaling in bacteria. Current cGAMP signaling pathways are found in deltaproteobacteria (red) and in the gammaproteobacterium *Vibrio cholerae* (blue). While the central second messenger, cGAMP, is identical, these pathways appear to use different classes of synthases, effectors, and phosphodiesterases. Text in red indicate new components revealed in this paper.

A distinguishing feature of cGAMP signaling is that two distinct pathways apparently arose from independent evolution in different bacteria. The initial discovery of cGAMP as a second messenger resulted from characterization of the synthase DncV in *V. cholerae* El Tor (6). This enzyme is structurally related to the OAS-like family of enzymes that includes cGAS, a mammalian innate immune sensor that produces mixed-linkage cGAMP (8). More recently, we discovered a different class of GMP-AMP cyclases (GACs) found in diverse deltaproteobacteria, including the environmental species *Geobacter sulfurreducens*, predatory species *Bdellovibrio bacteriovorus* (now within the class Oligoflexia), and social species *Myxococcus xanthus*. GACs are structurally related to the GGDEF family of diguanylate cyclases (DGCs), which are classically associated with cyclic di-GMP signaling, but instead have ‘Hypr’ GGDEF domains harboring specific variations in the active site that favor cGAMP production (9). We recently demonstrated that the Hypr GGDEF GSU1658 (*Gs*GacA) from *Geobacter sulfurreducens* regulates cGAMP but not cyclic di-GMP levels *in vivo*, resulting in a specific phenotype distinct from the biofilm-associated phenotype for the DGC EsnD (7).

While our focus to date has been on understanding the basis for cGAMP production by GACs with Hypr GGDEF domains (7, 9), the regulation of GAC activity is relatively unexplored (Fig. 1). Specifically, it was not known how the conserved inhibitory site (I-site) often found in GGDEF domains acts to regulate Hypr GGDEF enzymes. Also, no activator of bacterial cGAMP signaling has yet been identified. *Gs*GacA and its homologs in other bacteria contain receiver (Rec) domains, but the partner histidine kinases and their signal inputs have not been identified. Based on structural and *in vitro* evidence, *V. cholerae* DncV is proposed to be inhibited by folates (10), but this effect has not been tested *in vivo*. In any case, DncV does not appear to require activation by an exogenous factor.

To elucidate these missing parts of the Hypr-cGAMP signaling pathway and to broaden its scope to other bacteria, we have analyzed the system in the social bacterium *Myxococcus xanthus*. *M. xanthus* contains two GACs, MXAN_4463 (*Mx*GacA) and MXAN_2643 (*Mx*GacB), with Rec-GGDEF and GAF-GGDEF domain architectures, respectively (Fig. 2A). *Mx*GacA is homologous to *Gs*GacA and has an orphan receiver domain that lacks an associated histidine kinase. Aravind and colleagues first identified the GAF domain by bioinformatics and coined the name for its presence in c<u>G</u>MP-specific phosphodiesterases, *Anabaena* <u>a</u>denylate cyclases, and *Escherichia coli* <u>F</u>hlA (11). Since then, GAF domains have been shown to respond to a diverse set of small molecule signals, including cyclic nucleotides, amino acids, and metabolites, as well as redox and light (12–17). Our initial *in vitro* experiments with *Mx*GacB included measuring enzyme activity in the presence of ATP, GTP, and related analogs. Serendipitously, high activity was observed in the presence of millimolar levels of ATP, which had not been observed with *Gs*GacA, a Rec-GGDEF. This led to the hypothesis that an adenine-containing compound may be a physiological activator of *Mx*GacB (herein referred to as GacB).

**Figure 2.**
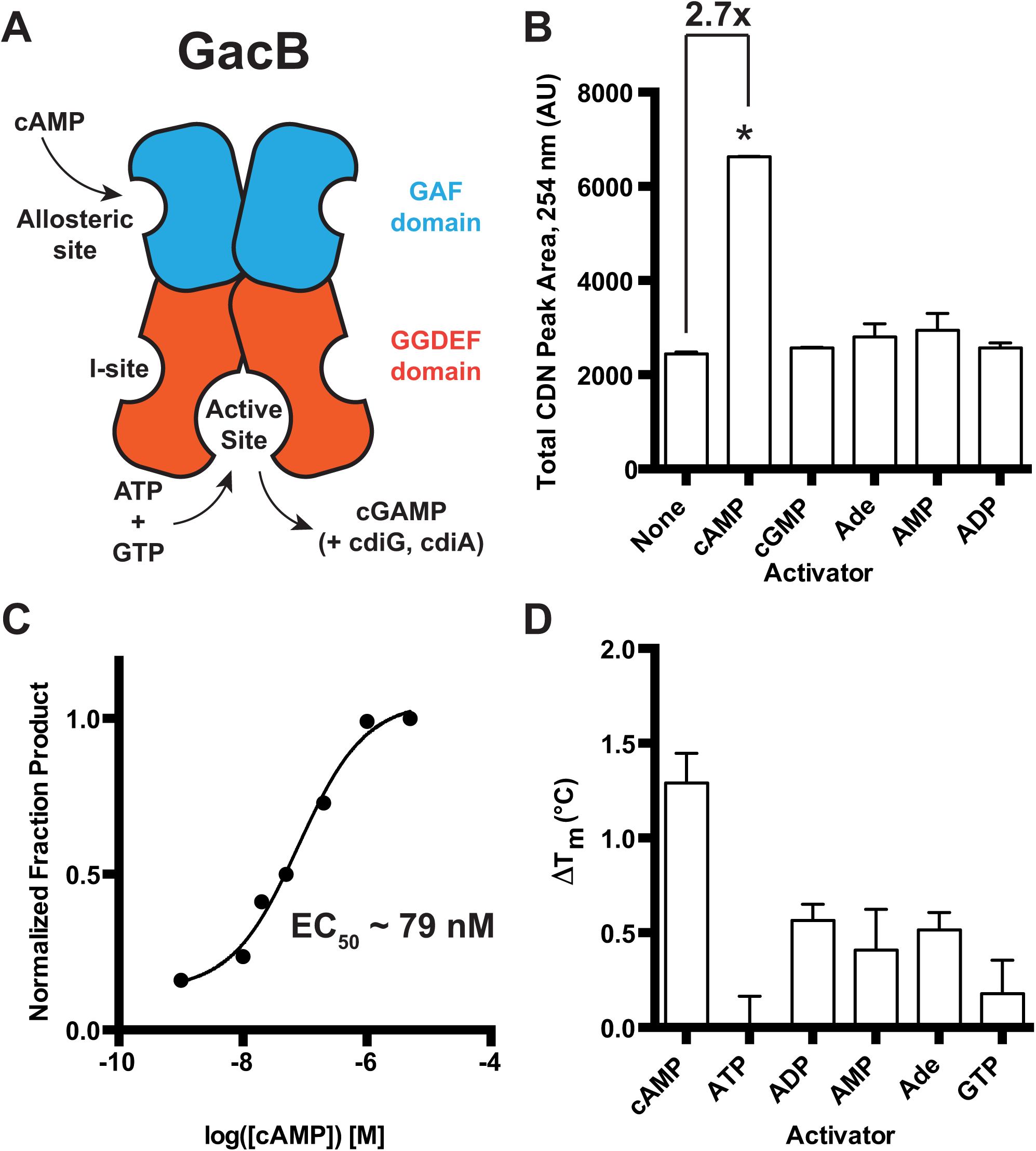
GacB, a Hypr GGDEF enzyme, is activated by cAMP. (A) Schematic of GacB showing the allosteric site on the N-terminal GAF domain (blue) and predicted inhibitory (I-site) in the GGDEF domain (red). (B) Synthase activity of GacB (0.5 μM) with 1:1 ATP and GTP was measured by LC-MS in the absence or presence of candidate activators (1 μM). The sum of cGAMP and cdiG UV peak areas at 254 nm is reported, as the amount of cdiA produced was undetectable (p <0.05, n=2). (C) Synthase activity of GacB (5 nM) with GTP doped with trace [α-^32^P] GTP was measured by TLC in the presence of increasing amounts of cAMP (0-5 μM). Production of cdiG was quantified by ^32^P signal density and plotted versus [cAMP] (n=1). (D) Ligand binding to the GacB GAF domain was assessed by thermal shift assay using an MBP-GAF domain construct (1 μM). ΔT_m_ was calculated as the difference between T_m_ with ligand (1 μM) and without ligand (n=3).

To identify the activator of GacB, an *in vitro* screen was conducted by incubating GacB with equimolar amounts of ATP and GTP substrates, with or without addition of candidate activators. A sum of all cyclic dinucleotide products was measured due to the promiscuous nature of GACs *in vitro*, an activity also seen with *Gs*GacA under these conditions (9). Only cAMP increased GacB activity (~2.7x) when added at a 10 μM concentration (Fig. 2B). Using a radiolabeled thin-layer chromatography (TLC) assay with GTP as the substrate to simplify product analysis, the effective concentration for GacB activation by cAMP (EC_50_) was found to be ~79 nM (Fig. 2C). A thermal shift assay conducted with the GAF domain of GacB (GacB_1-180_) shows that cAMP gives the greatest stabilization effect relative to other nucleotide ligands (Fig. 2D), supporting that the cyclic nucleotide binds in this domain. Taken together, these results demonstrate that cAMP is a selective activator that enhances GacB activity upon binding to the GAF domain.

GAF domains from bacteria and eukaryotes that bind cAMP or cGMP have been previously identified and structurally analyzed (12), including a recent structure of a cAMP-binding DGC enzyme, Lcd1 from *Leptospira interrogans* (18). However, to our knowledge, this is the first evidence linking cAMP signaling to cGAMP signaling, which expands the type of cAMP effectors in bacteria to include GAC enzymes. In addition, cAMP is the first identified small molecule activator of cGAMP synthesis, which provides insight into the physiological role of this signaling pathway in Myxobacteria, as GacB is conserved within the genus and in the related fruiting Myxobacteria *Corallococcus coralloides* and *Stigmatella aurantiaca* (19)

Besides regulation by sensory domains like the GAF domain, canonical GGDEF domain-containing DGCs also contain an allosteric inhibitory site (I-site) that binds cyclic di-GMP and leads to allosteric feedback inhibition of DGC activity (20). However, since GacB is a GMP-AMP cyclase with a divergent Hypr GGDEF domain, we had to consider whether it was selectively regulated by a given cyclic dinucleotide. GacB could be subject to product auto-inhibition by cGAMP, or it could be cross-regulated by cyclic di-GMP or cyclic di-AMP. Recent studies of *Gs*GacA yielded phenotypic and transcriptomic evidence consistent with cross-inhibition by cyclic di-GMP, but did not show a direct mechanism (7).

In our initial inhibition assays, both ATP and GTP were used as substrates, which complicated the analysis as GacB produced all three bacterial cyclic dinucleotides *in situ* (Fig. S1). We predicted that cyclic di-AMP was least likely to interact with GacB due to its structural dissimilarity to cyclic di-GMP, the canonical inhibitor of GGDEF enzymes. Therefore, the assays were repeated using ATP as the sole substrate, taking advantage of the *in vitro* promiscuity of GacB so that only cyclic di-AMP was produced *in situ*. Clear inhibition of enzyme activity was observed with 10 μM of cyclic di-GMP but with none of the other cyclic dinucleotides (Fig. 3A). We tested two potential mutants to knock out the I-site, GacB R286A and R292 (data not shown), and found that both had lower activity relative to WT, but R286A was not inhibited by cyclic dinucleotides. The IC_50_ value was determined to be 17.80 +/-1.06 μM for WT GacB inhibition by cyclic di-GMP (Fig. 3B), which is similar to the published IC_50_ value (5.1 +/-1.4 μM) for the DGC PleD by cyclic di-GMP (21).

**Figure 3.**
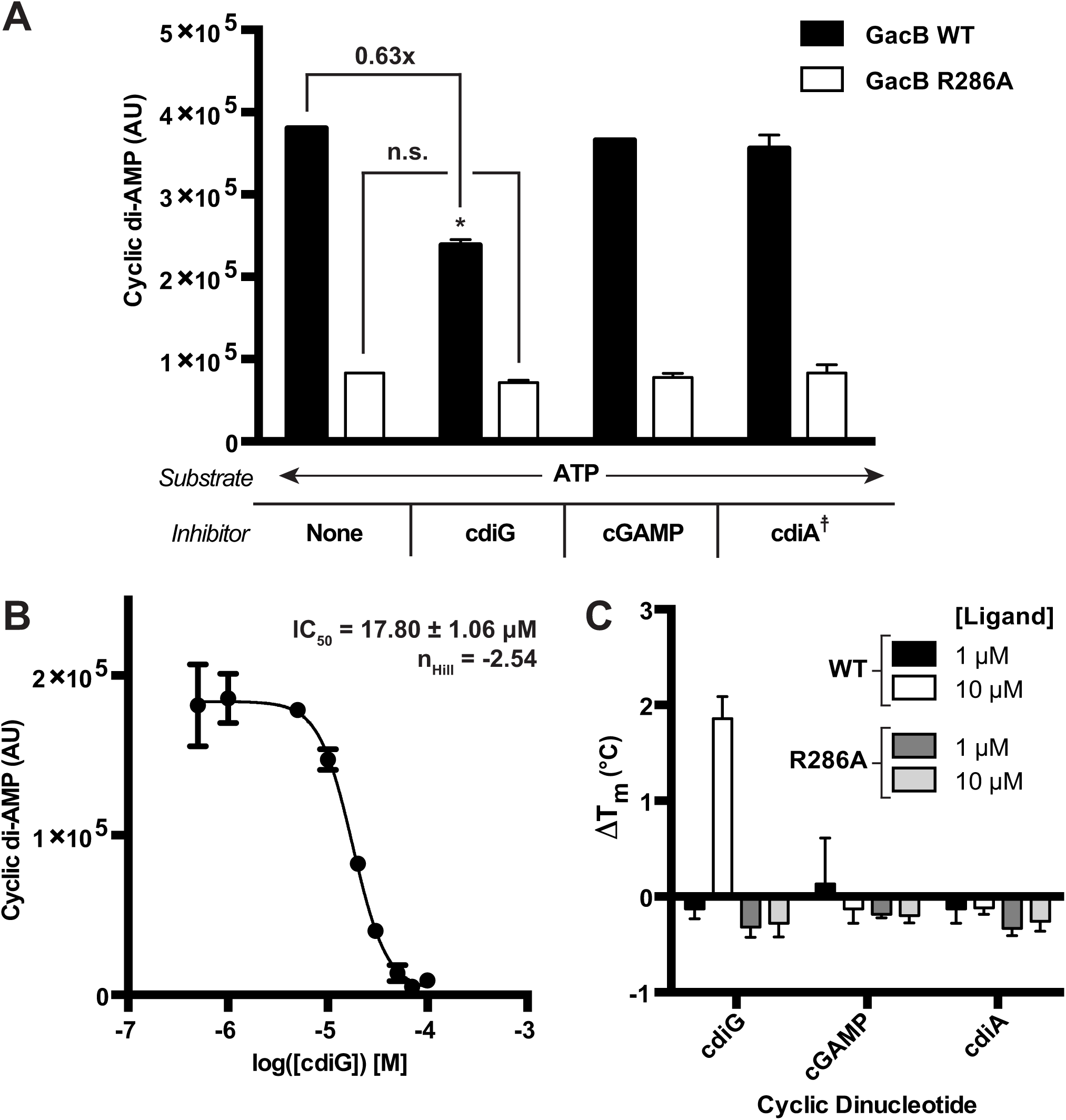
GacB is selectively inhibited by cdiG. (A) Inhibition of GacB WT or I-site mutant (1 μM) by CDNs (10 μM) was measured by LC-MS (p<0.05, n=2). Due to potential *in situ* inhibition of GacB by other CDNs, cdiA production from ATP as sole substrate was used to quantify activity. The product cdiA (659 *m/z*) is reported as an ion-extracted signal from the mass spectra. For reactions testing cdiA as the candidate inhibitor, the background signal from added cdiA was subtracted (╪). (B) Determination of IC_50_ value for cdiG inhibition of GacB using LC-MS (n=3). (C) CDN binding to full-length GacB WT or I-site mutant was assessed by thermal shift assay using MBP-GacB (1 μM) and 1 and 10 μM of each CDN. ΔT_m_ was calculated as the difference between T_m_ with CDN and without (n=3).

We also conducted a thermal shift assay of full length GacB WT and R286A with various amounts of each CDN. Significant stabilization of GacB WT was only observed with 10 μM of cyclic di-GMP, which is consistent with the activity assays and measured IC_50_ value (Fig. 3C). GacB R286A was not stabilized by cyclic dinucleotides. Thus, cyclic di-GMP binds and inhibits both Hypr and canonical GGDEF enzymes via the conserved I-site. However, whereas I-site regulation leads to auto-inhibition for DGCs, our data strongly support that inhibition of GacB by cyclic di-GMP instead is a mechanism for cross-regulation of cGAMP signaling by cyclic di-GMP.

Signal output can be decreased by inhibition of cGAMP synthesis or by degradation of cGAMP. So far, no cGAMP-specific PDE has been identified in bacteria. Three PDEs harboring HD-GYP domains from *V. cholerae*, including VC0681, were shown to degrade 3’,3’-cGAMP in preference to the mammalian mixed-linkage 2’,3’-cGAMP. When 3’,3’-linkage cyclic dinucleotides are compared, however, VC0681 clearly favors degradation of cyclic di-GMP over 3’,3’-cGAMP (22). VC0681 previously was shown to control cyclic di-GMP levels in *V. cholerae* (23), so does not appear to act as a cGAMP-specific PDE in cells.

We had hypothesized that the ‘erasers’ involved in cGAMP signaling would be specific, but still may have evolved from components of cyclic di-GMP signaling, in line with how we discovered GACs and cGAMP-specific riboswitches as sub-classes of GGDEF enzymes and GEMM-I riboswitches, respectively (9, 24, 25). In both cases, it was possible to identify specific residues in the active site or ligand binding pocket that rationally reprogrammed the synthase or riboswitch to gain new specificities. The *M. xanthus* genome contains six HD-GYP and two EAL domain containing genes that are candidate phosphodiesterases. Based on structure-guided sequence analysis, both *M. xanthus* EAL proteins contain conserved residues that enforce selectivity for cyclic di-GMP (26). However, when we compared to the HD-GYP domain from PmGH, a cyclic di-GMP PDE, we found that one *M. xanthus* HD-GYP protein does not harbor the two basic residues, R314 and K317, of the conserved Rxx(K/R) motif that recognizes a guanine nucleobase of cyclic di-GMP (27). This exception was MXAN_2061, also called PmxA, which contains an R515 and Q518 pair instead. *In vitro* studies identified PmxA as a cyclic di-GMP PDE, but deletion of this gene did not show significant changes in cyclic di-GMP levels *in vivo* (28). The switch from a hydrogen-bond donor (K/R) to a potential hydrogen-bond acceptor (Q) was promising for altering specificity toward cGAMP (Fig. 4).

**Figure 4.**
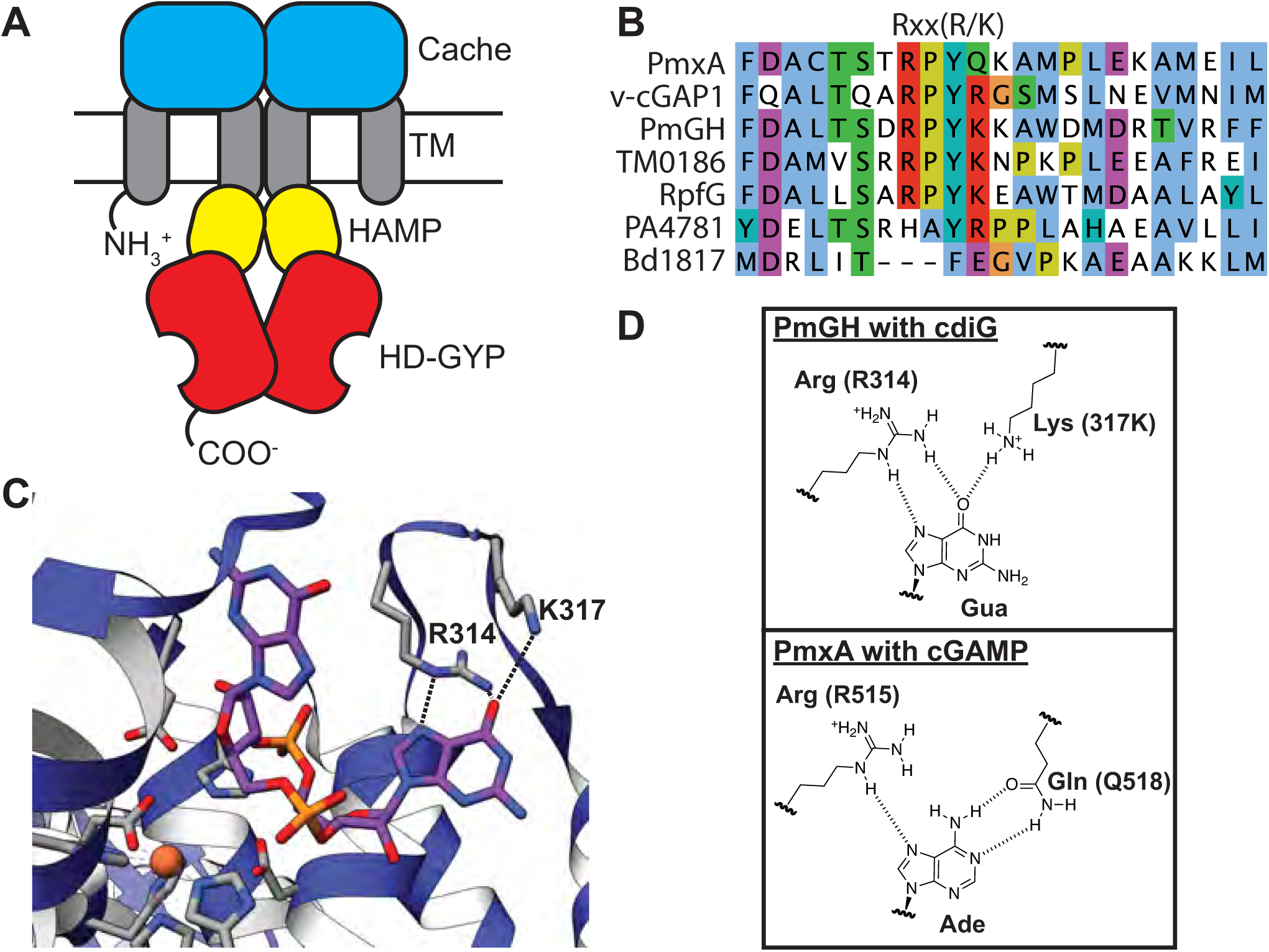
Sequence and structural analysis of PmxA reveals signature active site variation. (A) Schematic of the PmxA (MXAN_2061) predicted homodimer. A single monomer contains an N-terminal sensory Cache domain, two TM helices, a coiled-coiled HAMP domain, and C-terminal HD-GYP domain with PDE activity. (B) Sequence alignment of PmxA with previously reported HD-GYP domain PDEs using the MUSCLE alignment tool. The canonical Rxx(R/K) motif is shown. PmxA has a Gln (Q) at the fourth position of the motif. (C) View of the active site of cdiG PDE PmGH (PDB 4MDZ) from *Persephonella marina* shows the two canonical motif residues, R314 and K317, forming hydrogen bond contacts with one of the two guanine nucleobases in cdiG. (D) Molecular diagrams that show the structurally characterized interaction of the PmGH RxxK motif with cdiG and the proposed interaction of PmxA RxxQ motif with cGAMP.

PmxA is a multi-domain protein, composed of two transmembrane regions, a coiled-coiled HAMP domain, and the catalytic C-terminal HD-GYP domain (Fig. 4A). The Søgaard-Andersen group previously identified a truncated PmxA construct that contains the HD-GYP domain that was soluble and active, as well as developed buffer conditions to assay its PDE activity (28), which were critical advances for *in vitro* studies (Fig. 5A). Using an MBP-tagged version of this construct, we detected robust degradation of cGAMP after 4 h, whereas little to no degradation was observed for cyclic di-GMP and cyclic di-AMP under the same conditions (Fig. 5B). The AA-GYP double mutant rendered the PDE inactive toward cGAMP, confirming that the HD-GYP domain was necessary for this activity (Fig. S2).

**Figure 5.**
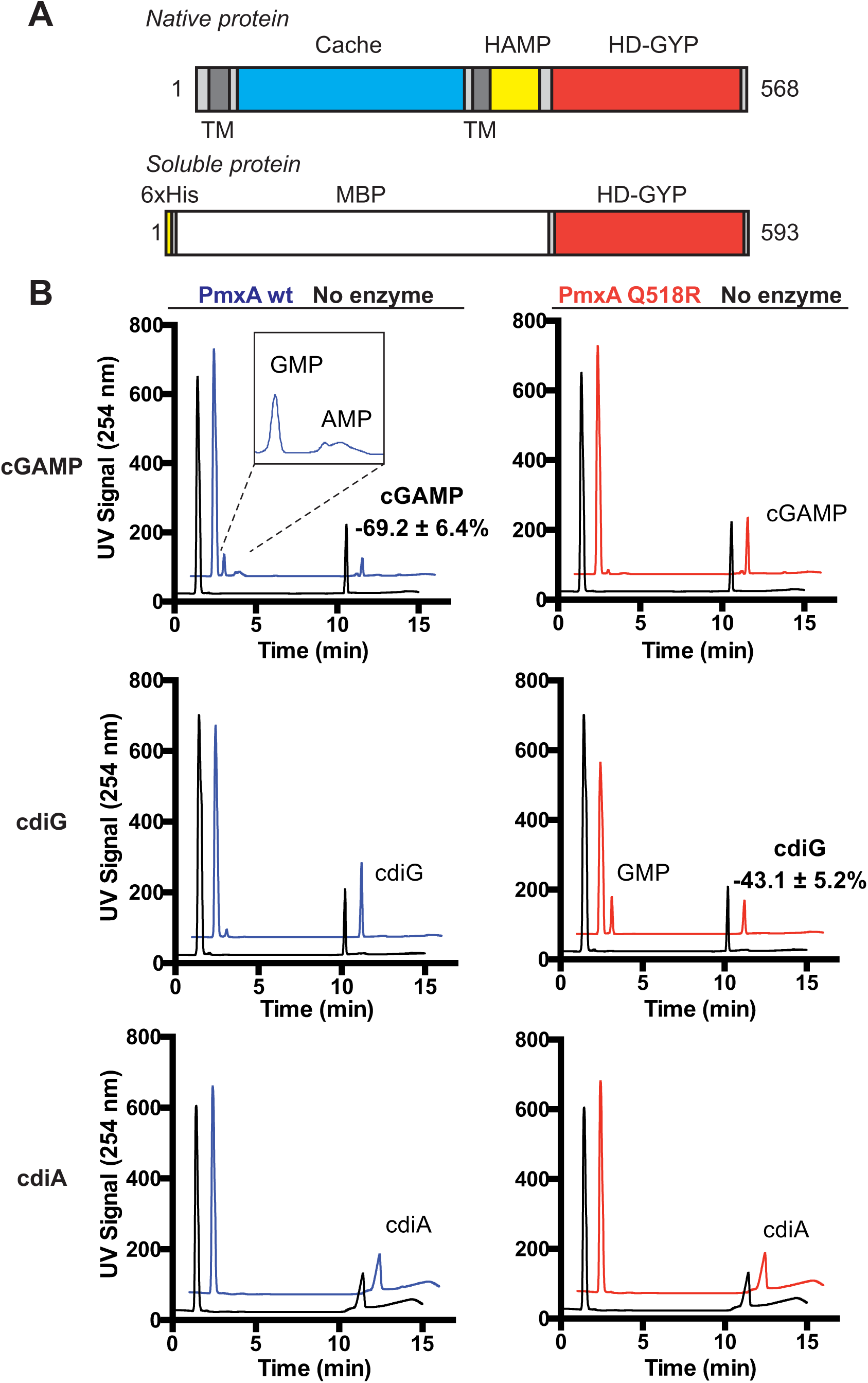
PmxA is a selective cGAMP phosphodiesterase that can be reprogrammed via the signature residue. (A) Schematic of native PmxA and the MBP-tagged soluble PmxA construct used for *in vitro* activity assays, which was first reported by Skotnicka *et al*. (B) *In vitro* PDE activity of MBP-PmxA wt (blue) and Q518R mutant (red) was analyzed by LC-MS. Enzyme (10 μM) or control (no enzyme, black) was incubated with 50 μM of cGAMP (top), cdiG (middle), or cdiA (bottom) for 4 h at 30 °C (n=2). Representative UV traces for each reaction are shown, with CDN peaks labeled based on MS analysis. Inset highlights the observation of mononucleotide products for PmxA wt with cGAMP.

To test whether the Q518 residue was responsible for the unique substrate selectivity of PmxA, the activity of the Q518R mutant PmxA was analyzed. This mutant exhibited little to no degradation of cGAMP after 4 h, and instead gained activity against cyclic di-GMP (Fig. 5B). The substrate selectivity of this point mutant is comparable to the published activity of v-cGAP1 (VC0681), which harbors an R residue at this position (22). To our knowledge, PmxA is the first bacterial PDE shown to preferentially degrade 3’,3’-cGAMP over cyclic di-GMP. Moreover, our data reveal that Q518 may serve as a signature residue for the sub-class of HD-GYP domains that act specifically as cyclic GMP-AMP phosphodiesterases (GAPs).

To investigate the regulatory function of cGAMP signaling in *M. xanthus*, the phenotype of a wild type strain (DZ2) was compared to the corresponding *ΔgacB* and *ΔpmxA* mutants. In a different strain, *M. xanthus* DK1622, the *ΔpmxA* mutant did not show a motility phenotype (19). Type IV pilus (T4P)-dependent motility, a.k.a. twitching or social motility, was assayed on CYE plates that contained 0.5% agar. Δ*gacB* slightly increased colony expansion (+22%) relative to WT, whereas Δ*pmxA* had the opposite effect (−24%) (Fig. 6A). These data indicate that cGAMP mildly inhibits T4P-dependent motility under these conditions.

**Figure 6.**
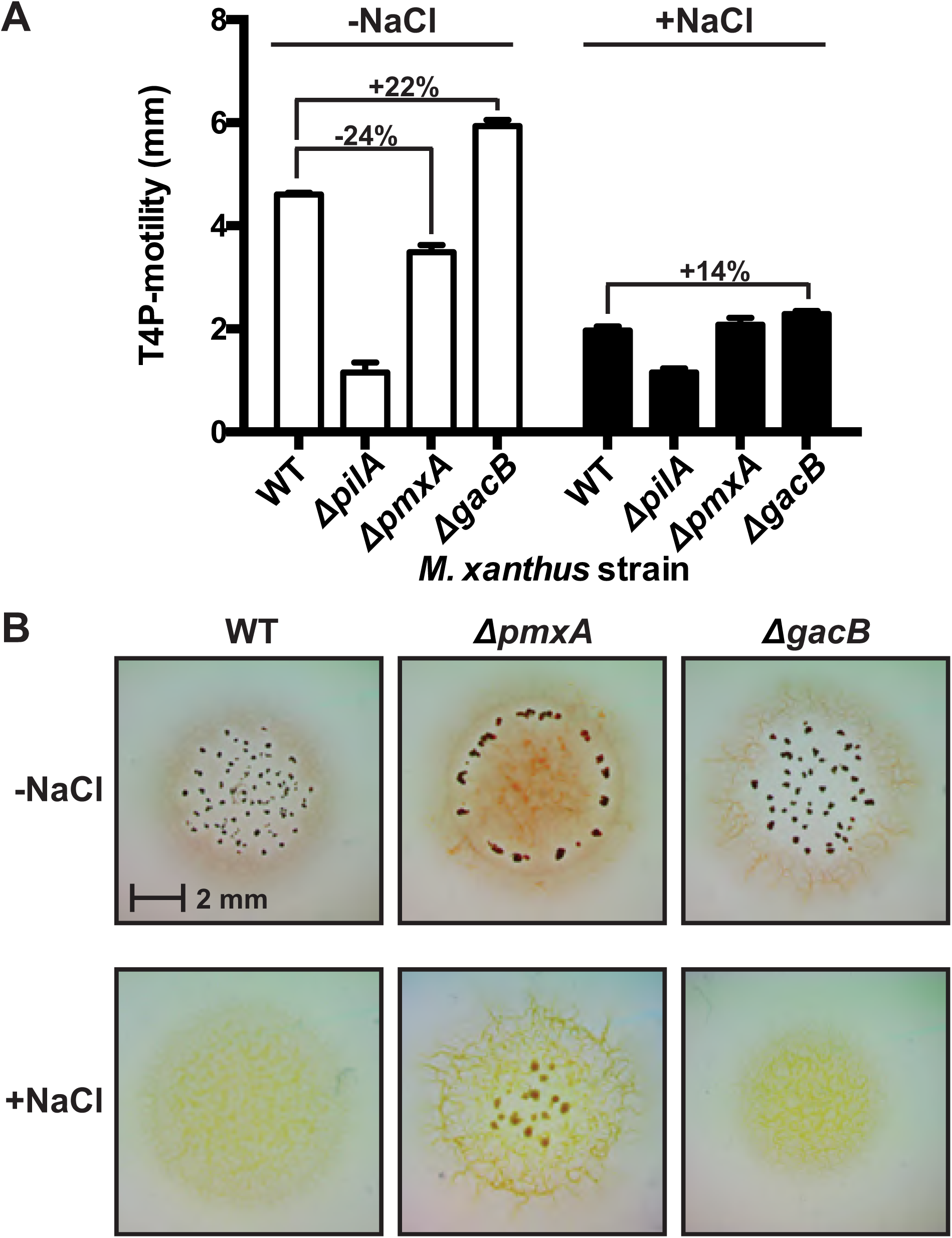
PmxA has mild effects on type IV motility but promotes resistance to osmotic stress in *M. xanthus*. (A) Type IV pili (S) motility is mildly inhibited by cGAMP in *M. xanthus*, as shown by comparison of wild-type strain (DZ2) versus Δ*pmxA* and Δ*gacB* mutants. This effect is less apparent under high osmolarity conditions (0.15 M NaCl, black), which globally suppresses T4P motility. (B) The Δ*pmxA* mutant forms fruiting bodies in high (+NaCl) or low (-NaCl) salt conditions, whereas fruiting body formation is inhibited in both wild-type (DZ2) and Δ*gacB* strains under osmotic stress conditions. Interestingly, the Δ*pmxA* strain forms fruiting bodies at the periphery of the swarm at low osmolarity and towards the center under high osmolarity.

Earlier studies reported that osmotic stress increases the concentration of cAMP in *M. xanthus* (29, 30). Since cAMP activates the synthase activity of GacB (Fig. 2B), we hypothesized that cGAMP may have a role during osmotic stress. Osmotic stress was induced by addition of high salt (0.15 M NaCl) to the growth medium. Under osmotic stress, we expected that cAMP stimulation of GacB activity should lead to higher cellular cGAMP concentrations that further inhibit T4P-dependent motility, whereas PmxA activity should have the opposite effect. However, we found that osmotic stress overall inhibits T4P-dependent motility in the wild type, *ΔgacB*, and *ΔpmxA* mutants (Fig. 6A). Consistent with the mild inhibitory phenotype observed for cGAMP, the *ΔgacB* strain that should produce less cGAMP than WT has slightly increased colony expansion (+14%) relative to WT, but the *ΔpmxA* mutant retained similar colony expansion compared to the wild type under high osmotic stress, even though it is predicted to have higher cGAMP levels. Taken together, our data suggest that cGAMP is not a major regulator of T4P-dependent motility in *M. xanthus*. These results contrast with cyclic di-GMP signaling in *M. xanthus*, which has been shown to strongly inhibit T4P-dependent motility (19), thus underscoring that cGAMP signaling is a distinct pathway from cyclic di-GMP in this organism.

Instead, we observe a developmental phenotype for cGAMP signaling that is apparent upon osmotic stress. *M. xanthus* form fruiting bodies to resist harsh environmental stresses and are induced, for example, under starvation conditions. When starved on CF agar in the absence of osmotic stress, all three tested strains (WT, Δ*pmxA*, and Δ*gacB*) formed fruiting bodies, albeit the fruiting bodies formed by the Δ*pmxA* strain are more peripheral. In contrast, in the presence of osmotic stress, only Δ*pmxA* is capable of forming fruiting bodies (Fig. 6B). This striking result indicates that high cGAMP levels provide resistance against osmotic stress during *M. xanthus* fruiting body formation. In a different strain, *M. xanthus* DK1622, the *ΔpmxA* mutant was shown to exhibit developmental defects, including irregular, translucent fruiting body formation and sporulation at 15% of WT levels without osmotic stress (28).

To further explore the function of PmxA and its homologs in cells, we developed a fluorescent biosensor assay for PDE activity based on RNA-based fluorescent (RBF) biosensors. RBF biosensors specific for each of the bacterial cyclic dinucleotides have been used previously to analyze synthase activities (31). This assay was adapted to screen PDE activities by encoding a candidate PDE in one plasmid and a matched pair of biosensor and synthase on a second plasmid (Fig. 7A). For example, the DGC WspR was co-expressed with the Dp biosensor, which responds to cyclic di-GMP (32), and the well-characterized GAC *Gs*GacA was co-expressed with the Gm biosensor, which responds to cGAMP (24). These two matched synthase-sensor pairs result in highly fluorescent (bright) cell populations as measured by flow cytometry. Co-expression of an active PDE then should cause the bright cell population to decrease, due to degradation of the cyclic dinucleotide sensed by the RBF biosensor. Thus, PDE selectivity can be preliminarily assessed by screening against the two synthase-sensor pairs.

**Figure 7.**
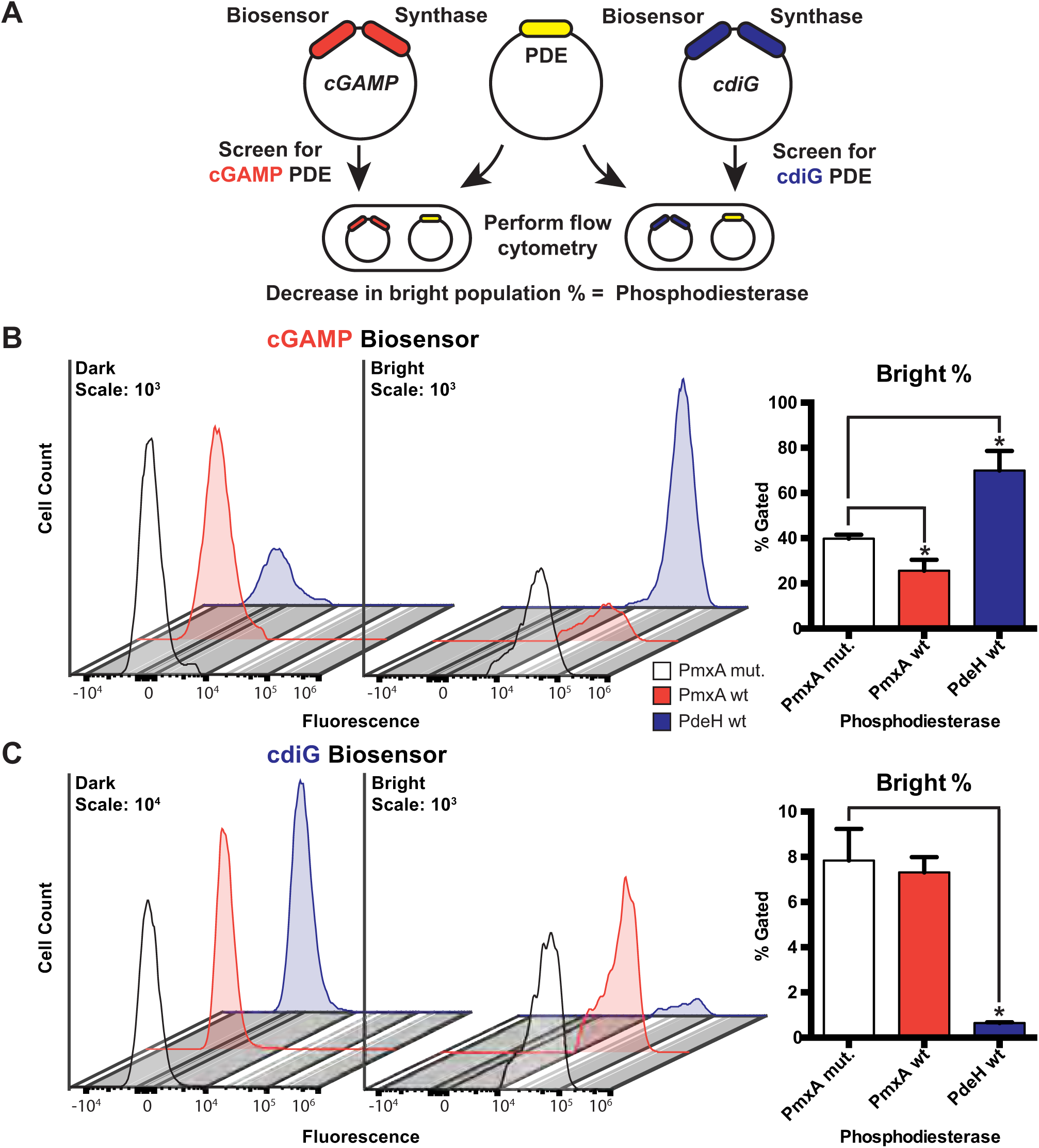
An RNA-based fluorescent biosensor assay for cyclic dinucleotide phosphodiesterases. (A) Schematic of RBF biosensor assay showing that a PDE candidate (yellow) and cGAMP (red) or cdiG (blue) biosensor-synthase plasmids are co-transformed and overexpressed in *E. coli* BL21* (DE3) cells. Active PDEs lead to a decrease in the bright cell population as analyzed by flow cytometry in the presence of the DFHBI dye. (B) Change in cellular cGAMP levels due to enzyme activity was measured by flow cytometry. Representative histograms show the dark (left) and bright (right) cell populations upon expression of WT PmxA (red), cdiG PDE PdeH (blue), and inactive PmxA mutant as a control (white). Y-axes are scaled by total cells (10^3^), while the x-axes depict logarithmic fluorescence intensity. Bar graph depicts the change in % bright relative to the inactive control (p<0.05, n=3). (C) Change in cellular cdiG levels due to enzyme activity was measured by flow cytometry. Same as in part B (p<0.05, n=3), except the histogram scale for the dark cell population is 10^4^.

Since this assay requires balanced co-expression of three components (synthase, biosensor, and PDE), growth and induction conditions had to be optimized, especially for a high-throughput, 96-well format. Two factors that significantly improved assay results were the culture volume per well and % lactose in the autoinduction media. We also found that the addition of a third component, even an inactive protein (PmxA mut, Fig. S2), leads to a mixed population of bright and dark cells rather than a single bright population as seen in prior studies co-expressing just synthase and biosensor (Fig. 7B, C). Most likely, this effect is due to stochastically reduced synthase activity from non-uniform expression levels. The synthases are particularly sensitive to this, because as GGDEF enzymes they are active only as dimers or higher order oligomers.

To assess PDE activity in the context of these bifurcated cell populations, we decided to analyze the percentage of the bright cells in the sample. Using this approach, it was verified that wild-type PmxA decreases the population of cGAMP-bright cells relative to the inactive PmxA mutant, but it has no effect on the population of cyclic di-GMP-bright cells (Fig. 7B, C). This result shows that PmxA is a cGAMP-specific PDE in the live cell context and corroborates our biochemical data.

We had expected the opposite trend upon addition of PdeH, which is a well-studied cyclic di-GMP-specific PDE from *E. coli* previously called YhjH (33, 34). While PdeH did decrease the population of cyclic di-GMP-bright cells (Fig. 7C), it counterintuitively increased the population of cGAMP-bright cells. One reasonable explanation for this effect is that PdeH de-represses the *Gs*GacA synthase by lowering global cyclic di-GMP levels in *E. coli*. This result is in line with our demonstration that GAC activity is inhibited by cyclic di-GMP *in vitro* (Fig. 3), is consistent with reporter assay results in *G. sulfurreducens* (9), and further supports that this cross-regulation occurs in cells.

With this assay and analysis method validated using PmxA and PdeH, it was applied to screen HD-GYP domains from other bacteria for cGAMP-specific PDE (GAP) activity. We first performed a bioinformatic search outside the *Myxococcus* genus for HD-GYP domains homologous to PmxA that harbor the RxxQ motif, which was shown to be important for cGAMP specificity. HD-GYP domains with the RxxN motif were added to this search, as N contains an amide functional group similar to Q. The majority of homologs (87.97%) harbored the canonical Rxx(K/R) motif, but 539 sequences (4.69%) contained an Rxx(Q/N) sequence (Fig. 8A). Excitingly, while some candidates are from bacterial species previously predicted to have cGAMP signaling due to the presence of the synthase (9), the majority are from bacterial species not previously associated with cGAMP signaling (Fig. 8B). Candidate GAPs appear well represented in both Proteobacteria and Firmicutes.

**Figure 8.**
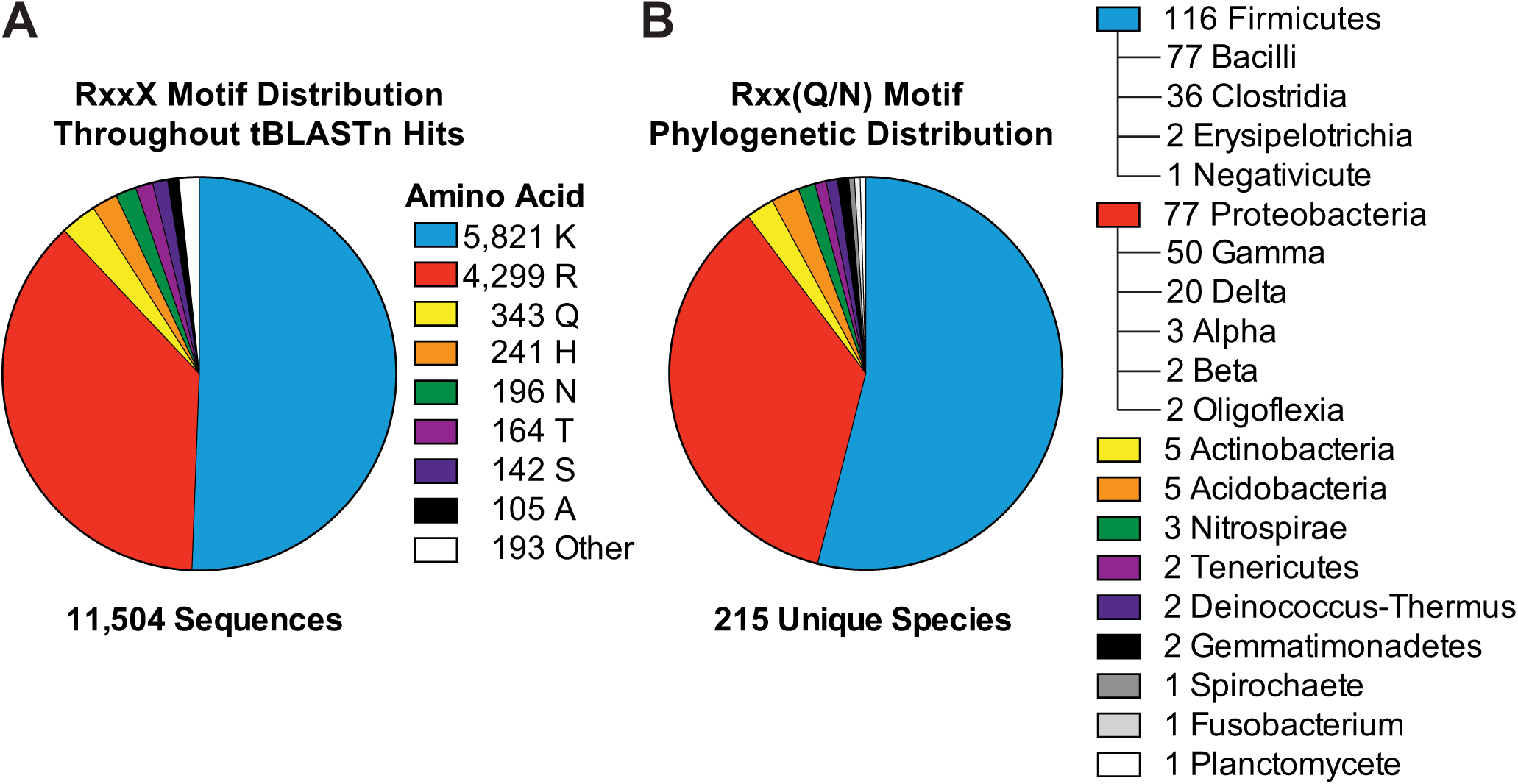
The variant signature Rxx(Q/N) motif is broadly distributed and most prevalent in Firmicutes and Proteobacteria. (A) The prevalence of the top 8 variants for the RxxX motif in bacterial HD-GYP domains is shown. The sequence analysis used 11,504 putative HD-GYP domains containing aligned RxxX motifs out of 14,326 total tBLASTn hits against the PmxA HD-GYP domain. (B) The distribution of RxxQ and RxxN HD-GYP motifs in Bacteria. A total of 539 sequences containing RxxQ or RxxN motifs were sorted based on unique species, leading to the identification of unique 215 bacterial species containing HD-GYP domains with these variant motifs. The species were categorized using NCBI Taxonomy.

Out of 539 total candidate GAPs matching the Rxx(Q/N) motif (RxxQ, 343; RxxN, 196), six candidate genes were selected based on their phylogenetic diversity. The full length PDEs were cloned and screened for activity against cGAMP and cyclic di-GMP using the *in vivo* biosensor assay (Fig. 8A). Gratifyingly, five out of six candidates showed PDE activity against cGAMP, whereas the last candidate appears inactive or poorly expressed in the assay. Of the five active PDEs, three appear selective for cGAMP and two appear to degrade both cyclic dinucleotides. Thus, the candidates from *Geobacter metallireducens, Bdellovibrio bacteriovorus*, and *Vibrio tapetis* are putative GAP enzymes, whereas additional analysis is required to determine substrate preferences for the enzymes from *Moorella thermoacetica* and *Phaeobacter piscinae*.

*B. bacteriovorus* HD-GYP Bd2325 (Bd) demonstrated the strongest cGAMP-specific PDE activity in our screen and harbors the alternative RxxN motif. Bd2325 was cloned as a full-length, N-terminal MBP-tagged protein for testing *in vitro*. No PDE activity against either cGAMP or cyclic di-GMP was observed using *in vitro* assay conditions developed for PmxA, which included 10 mM Mg^2+^ as the added divalent metal (Fig. 9A, B). However, different divalent metals and pH have been shown to enhance HD-GYP activity (22, 35), as HD-GYP active sites are known to contain redox-sensitive bi-and tri-nuclear cores with variable metal selectivity (36–38). We screened Bd2325 with different metals (Mg^2+^, Mn^2+^, Ca^2+^, Fe^2+^, and Ni^2+^, Fig. S3) and pH ranges (data not shown), and indeed found that two types of divalent metals activate the enzyme *in vitro*.

**Figure 9.**
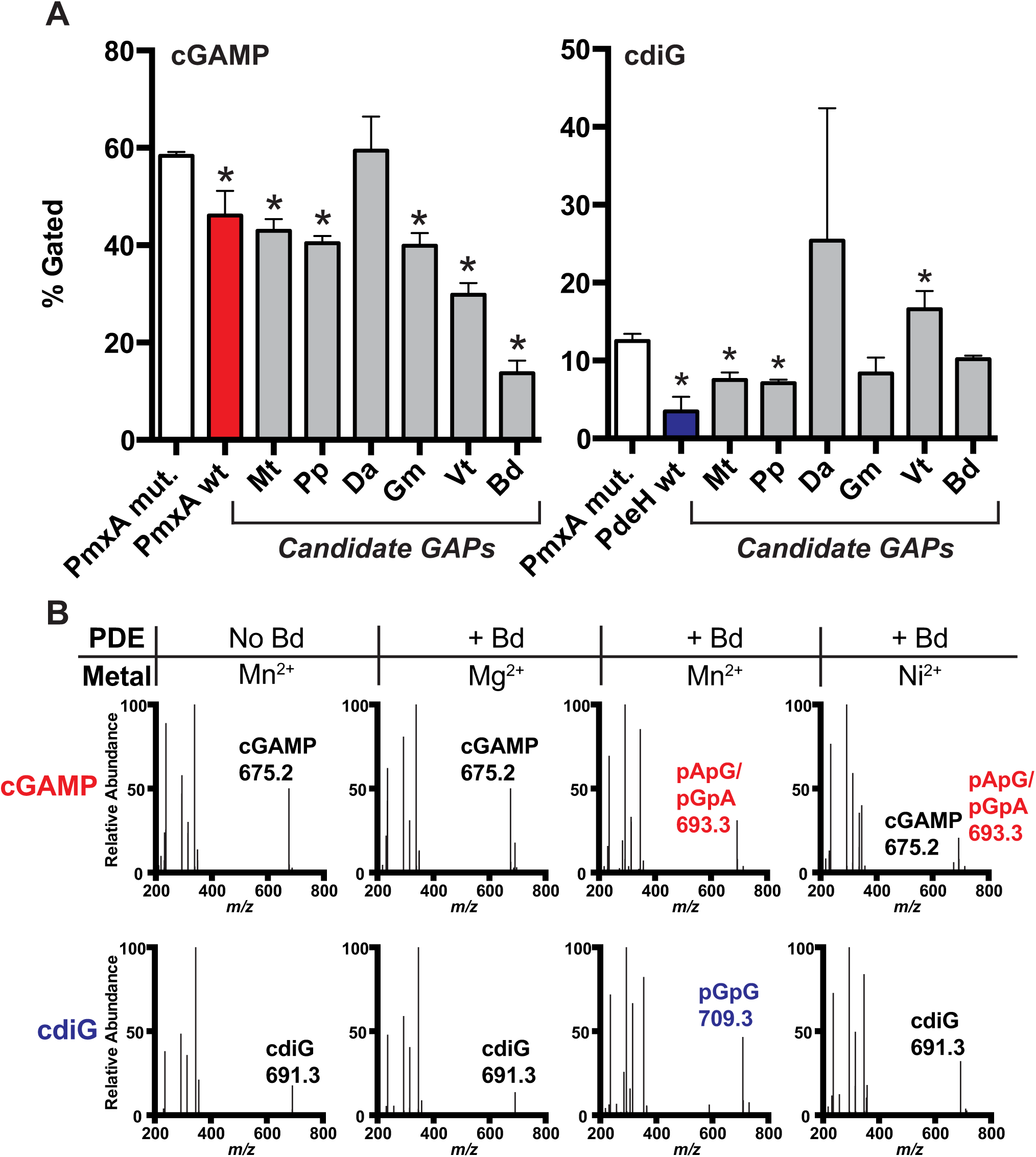
Identification of selective cGAMP phosphodiesterases (GAPs) from other bacteria, including one with the RxxN motif. (A) PDE activity screen of six candidate GAPs using the biosensor-based flow cytometry assay depicted in **Fig. 7A**. PDEs contain a Q (Mt: *Moorella thermoacetica*, Pp: *Phaeobacter piscinae*, Da: *Dechloromonas 19romatic*, Gm: *Geobacter metallireducens*, Vt: *Vibrio tapetis*) or N (Bd: *Bdellovibrio bacteriovorus*) at the selectivity position. Activity of PDE candidates were compared to an inactive control, PmxA mutant (p < 0.05, n=3). (B) *In vitro* PDE activity of MBP-Bd2325 with different divalent metals was analyzed by LC-MS. Enzyme (10 μM) or control (no enzyme) was incubated with 50 μM of cGAMP or cdiG for 4 h at 30 °C in the presence of Mg^2+^, Mn^2+^, or Ni^2+^. A representative mass spectrum for each reaction is shown, with CDN or product peaks labeled based on MS analysis (n=2).

Bd2325 is most active in the presence of Mn^2+^, as it degraded both cGAMP and cyclic di-GMP to their respective linear dinucleotides within 4 h (Fig. 9A, B). In fact, under these conditions, this enzyme degraded a measurable amount of cGAMP within the ~30 seconds required to prepare the ‘0’ minute time point of a time course, but was less active against cyclic di-GMP (Fig. S4). Observation of the linear dinucleotide product contrasts with the mononucleotides AMP and GMP detected with PmxA. HD-GYP protein TM0186 from *Thermotoga maritima* shifts from production of linear to mononucleotide product depending on active site metal occupancy, which may also be true for other HD-GYP PDEs (38). The preference for cGAMP was even clearer in the presence of Ni^2+^. After 4 h, cGAMP was fully degraded, but the majority of the cyclic di-GMP sample remained. Thus, these *in vitro* experiments confirm that variant HD-GYP domains harboring either the RxxQ or RxxN motifs are selective for cGAMP over cyclic di-GMP.

According to our bioinformatics analysis, the newfound GAP enzymes are more widespread in bacteria than the known set of Hypr GGDEF synthases and cGAMP riboswitches that have defined the Hypr-cGAMP signaling pathway to date. Thus, it is plausible that they also play a role in the DncV-cGAMP signaling pathway, although the previously described v-cGAPs in *V. cholerae* do not harbor the active site variations that give rise to cGAMP specificity. Going forward, it will be interesting to determine whether these two cGAMP signaling pathways have points of convergence, like utilizing the same PDE class, or will remain completely distinct from one another in terms of signaling components. A further intriguing possibility we want to pursue is that some HD-GYP variants may have activity against newly discovered cyclic dinucleotide signals (39). There is still high potential for novel enzyme classes associated with cyclic dinucleotide signaling, as well as novel effectors that bind cyclic dinucleotides.

When cellular signaling pathways are highly integrated, phenotype-based studies may become challenging to interpret or use for deconvoluting pathways and discovering novel components. To address this issue, we have worked on developing high-throughput fluorescence assays that read out specific cyclic dinucleotide levels in vivo (31). We recently showed that RBF biosensors can be applied to detect modulation of endogenous cyclic di-GMP signaling (40, 41). Here we show for the first time that RBF biosensors can be used to discover and evaluate specific PDE activity.

Taken together, this study expands our knowledge of how cGAMP signaling is regulated, by identifying the first activator molecule, cAMP, and the founding members of a class of cGAMP specific phosphodiesterases (GAPs). In addition, we show that cGAMP signaling is integrated with other nucleotide signaling pathways via allosteric regulation of the synthase, GacB, by cAMP and cyclic di-GMP. This result reinforces the growing appreciation that nucleotide signaling pathways form an integrated, multi-layered network, allowing bacterial cells to respond in a coordinated, sophisticated manner to complex feedback from their environment and metabolic status.

## MATERIALS AND METHODS

### In Vitro Activity Assay for GacB

For activation assays, 6xHis-MBP GacB WT (0.5 μM) was incubated in a 50 μL solution of 50 mM Tris-HCl pH 7.5, 10 mM MgCl_2_, 100 mM NaCl, and 5 mM dithiothreitol, with both substrates (ATP and GTP) at 1 mM each and 1 μM of candidate activators (cAMP, cGMP, adenosine, AMP, ADP, or none). Reaction mixtures were incubated at 37 °C for 6 h, heated to 95 °C for 30 s to denature the protein, then centrifuged for 3 min at 13,000 rpm to remove precipitate. For LC-MS analysis, 20 μL of the sample was injected, and starting materials and products were confirmed by mass and quantified by peak area at 254 nm absorbance.

Inhibition assays were performed following the same protocol as above, with the following changes. 6xHis-MBP GacB WT (0.5 μM) was first incubated in reaction buffer with 10 μM of candidate inhibitors (cGAMP, cdiG, cdiA, or none) for 10 min at room temperature. Then substrate (1 mM ATP) was added to start the reaction. Signal was measured by ion extraction of respective CDN m/z values: 659 m/z cdiA, 675 m/z cGAMP, 691 m/z cdiG. The IC_50_ value for GacB was obtained using this protocol, but with varied amounts of cdiG between 0-100 μM.

### Activity Assay for Phosphodiesterases using LC-MS

The protocol was adapted from previously published methods for PmxA developed by Skotnicka *et al.* (28). Briefly, enzyme (10 μM final concentration) was incubated at 30 °C for 5 min in buffer containing 50 mM Tris-HCl pH 8.0, 300 mM NaCl, and 10 mM MgCl_2_. Then, 50 μM substrate (cGAMP, cdiG, or cdiA) was added and the mixture was incubated at 30 °C for 4 h. Reactions were stopped by addition of an equal volume of 0.5 M EDTA, pH 8.0. Samples were incubated at 95 °C for 30 s to denature the enzyme, centrifuged for 3 min at 13,000 rpm to remove precipitate, and 20 μL of the sample was analyzed by LC-MS. Peak assignments were made based on *m/z* values. For experiments testing enzyme activity in presence of other divalent metals, 10 mM of each metal (MnCl_2_, CaCl_2_, FeSO_4_, NiSO_4_) was substituted for MgCl_2_ in the reaction buffer.

### Fluorescent Biosensor Screening Assay for Phosphodiesterases

Chemically competent *E. coli* BL21 (DE3) Star cells (Life Technologies) were co-transformed with different combinations of biosensor-synthase plasmid (pET-Duet with synthase in multicloning site 1 and biosensor in multicloning site 2) and candidate PDE plasmid (pCOLA-Duet with PDE gene cloned into multicloning site 2). The screen for cGAMP PDE activity used biosensor Gm790P1-4ΔA-Spinach2 (24) and synthase GacA from *G. sulfurreducens* (9). The screen for cyclic di-GMP PDE activity used biosensor Dp17-Spinach2 (32) and synthase WspR from *P. fluorescens* (42). Single colonies from LB/Carb/Kan plates (50 μg/mL each antibiotic) were used to inoculate 500 μL starter cultures in non-inducing media (43) with antibiotics in a 96-deep well culture plate (VWR). The 96-well starter cultures were grown 16-20 h at 37 °C with vigorous shaking at 325 rpm. After non-inducing growth, a 1:100 dilution of starter culture was added to ZYP autoinduction media modified to contain 1% α-lactose, which was found to improve expression and fluorescence signal. Culture volumes at a ratio of ~1:7 media:headspace (300 μL media in 2200 μL wells) were found to be critical for efficient induction of expression. Cells were grown under inducing conditions for 16-20 h at 37 °C with vigorous shaking at 325 rpm, then diluted 1:70 in 1x PBS, pH 7.4 with 50 μM DFHBI-1T for analysis by flow cytometry. Cellular fluorescence was measured for 50,000 cells at a flow rate of 25 μL/min using the Attune NxT Flow Cytometer (CIRM/QB3 Facility, UC Berkeley) on the BL1 channel (Ex. 530 +/-30 nm, PMT V=540). Data was analyzed with FlowJo software (version 10.5.3). Successive polygon gates were applied to isolate single *E. coli* cells. A final bisecting gate was applied to separate cells into bright versus dark populations. This bisecting gate isolated the individual bright (Mean Fluorescence Intensity ~55,000 AU) or dark (Mean Fluorescence Intensity ~1,500 AU) peak in the fluorescence histogram and was uniformly applied across all replicates for a given CDN tested. Results are reported as both representative fluorescence histograms and averages of % cells in the bright population.

### *Myxococcus xanthus* Phenotypic Assays

To construct the in-frame deletion strains, in-frame deletion mutation cassettes were amplified with polymerase chain reaction (PCR) using chromosomal DNA as template, digested and inserted into plasmid pBJ113. All constructs were confirmed by DNA sequencing. To generate mutant strains, transformants were obtained by homologous recombination as previously described (44), using electroporation of *M. xanthus* DZ2 cells with 4 μg of plasmid DNA. Mutants were confirmed by PCR and sequencing of the gene of interest. T4P-dependent motility and fruiting body formation were assayed on CYE plates containing 0.5% (w/v) agar and CF plates containing 1.5% (w/v) agar as described, respectively (44, 45). For T4P-dependent motility, colony expansion was determined as the increase in colony diameter after 24 h incubation at 32 °C. For fruiting body formation, cells were incubated at 32 °C for 72 h and 120 h under low (0 M NaCl) and high (0.15 M NaCl) osmolarity, respectively. A Nikon SMZ1000 microscope and an OMAX A3590U digital camera were used to capture the images of fruiting bodies.

*Additional data are available in Supplementary Information*

## ACKNOWLEDGEMENTS

The authors would like to acknowledge Dr. Mary West of the CIRM/QB3 Shared Stem Cell Facility at UC Berkeley for use of the Attune flow cytometer. We thank Dr. Wenjun Zhang and her lab for allowing TAW to share lab space at UC Berkeley, as well as Andrew Dippel for helpful discussion of cyclic dinucleotide signaling proteins. This work was supported in part by NSF grants 1716256 and 1915466 to MCH, NIH grants R01 GM124589 (to MCH) and GM129000 (to BN), and UC Berkeley College of Chemistry undergraduate summer research awards (to JJP, LJ, and WAA).

