## Supplementary Information for "Second messengers and divergent HD-GYP enzymes regulate 3’,3’-cGAMP signaling"

**This PDF file includes:**

Extended Materials and Methods
Figs. S1 to S4
Tables S1 to S2
References for SI reference citations

### Extended Material and Methods

#### General Reagents and Oligonucleotides

Oligonucleotide primers were purchased from either Elim Biopharmaceuticals (Hayward, CA) or Integrated DNA Technologies (Coralville, IA). *M. xanthus* genes were amplified from genomic DNA. GacB mutant R292A was previously reported in Hallberg et al. (1). Candidate cGAMP PDE genes were purchased as gBlocks (IDT). Nucleotide reagents for activator screening were obtained from Sigma-Aldrich (St. Louis, MO), Acros Organics (Belgium), and Biolog (Germany). Cyclic dinucleotide standards for inhibition assays and phosphodiesterase experiments were purchased from Biolog and Axxora (Farmingdale, NY). PEI-cellulose TLC plates and Sypro Orange (5,000x in DMSO) were bought from Sigma-Aldrich. DFHBI-1T used for flow cytometry screening was chemically synthesized from established protocols (2).

#### Molecular Cloning

All gene constructs for *in vitro* assays were cloned into a pET16 vector containing an N-terminal purification tag that included both 6xHis tag and maltose binding protein (MBP) between NdeI and BamHI restriction sites. Gene constructs were cloned as fusions to 6xHis-MBP using BamHI and NotI restriction sites. PmxA was analyzed as the truncation PmxA$^{384\text{-}568}$ containing the HD-GYP domain first described by Skotnicka *et al.* (3). Mutations were generated by the around-the-horn method. Untagged candidate PDE genes for flow cytometry assays were cloned into pCOLADuet-1 between restriction sites NdeI and XhoI in the second multi-cloning site. Alternative sites were used for sequences where these cut sites were found within the gene (BglII/XhoI for Pp, NdeI/KpnI for Gm). Gibson assembly was used to construct vectors containing a CDN synthase and biosensor in plasmid pETDuet-1. The synthase (WspR or *Gs*GacA) was placed 5' to the biosensor (Dp17 or Gm790P1-4ΔA) under control of separate T7 promoters, with a short spacer sequence in between. Appropriate overhangs for Gibson assembly were added to the pETDuet-1 vector backbone, the synthase gene (WspR or *Gs*GacA), and the biosensors (Dp or Gm) by PCR with extended primers (Table S1). Templates for the biosensor have the design T7 promoter – $tRNA_{Lys}$(5' half)– Spinach2(5' half) – biosensor – Spinach2(3' half) – $tRNA_{Lys}$(3' half)– T7 terminator. These components were combined through a three piece Gibson assembly into the final plasmid construct.

#### Protein Overexpression and Purification

Proteins for *in vitro* analysis were overexpressed in BL21* (DE3) cells using either ZYP-5052 autoinduction media (4) or 2x YT media with IPTG induction with carbenicillin at 50 µg/mL final concentration. Autoinduction was used for production of GacB and PmxA constructs, whereas IPTG induction was used for Bd2325 due to poor expression observed in ZYP-5052 media. In ZYP-5052 media, cultures were inoculated and grown at 37 °C for 20-24 h. In 2x YT media, cells were grown at 37 °C until $OD_{600}$ ~ 0.6-0.8 and induced with 1 mM IPTG, followed by 20 h expression at 18 °C. Cultures were grown at either 250 or 750 mL volumes. Subsequent purification steps were identical for all proteins. Cells were harvested by centrifugation and lysed by sonication in lysis buffer containing 25 mM Tris-HCl pH 8.2, 500 mM NaCl, 30 mM imidazole, 5 mM 2-mercaptoethanol, and 5% glycerol. Clarified lysates were generated by centrifugation at 9,200 rpm for 40 min then were incubated with 1.5-4 mL HisPur™ Ni-NTA Resin (Thermo Scientific). The resin was loaded onto a disposable protein chromatography column, the flow-through was drained, and the resin bed was washed with 60 mL lysis buffer. Protein was eluted with 10 mL lysis buffer containing 300 mM imidazole. Eluted proteins were buffer exchanged by dialysis or centrifugation into storage buffer containing 20 mM HEPES pH 7.5, 250 mM KCl, 5 mM 2-mercaptoethanol, and 5% glycerol, followed by concentration using centrifugal concentrator columns. Final protein concentrations were calculated by absorbance at 280 nm. Purified protein aliquots were flash frozen in liquid nitrogen and stored at -80 ºC until further use.

**LC-MS Analysis of In Vitro Enzyme Activity Assays**

LC-MS analysis of in vitro enzyme activity assays was performed on an Agilent 6120 Quadrupole MS with an Agilent 1260 Infinity HPLC equipped with a multi-wavelength detector (MWD). Samples were separated on a Poroshell 120 EC C18 column (50 mm length x 4.6 mm internal diameter, 2.7 μm particle size, Agilent) at a flow rate of 0.4 mL/min. The solvent system (aqueous 10 mM $NH_4OAc$, 0.1% AcOH as Solvent A and MeOH as Solvent B) and gradient was based on the procedure developed by Burhenne *et al.* (5). Under these conditions, cyclic dinucleotides eluted between 10-12 min with an elution peak order of cdiG, cGAMP, and cdiA observed at 254 nm. For phosphodiesterase assays, linear products eluted in a similar range of retention times. Molecular assignment was made through analysis of the mass spectra. MS was conducted in positive ion mode with an electrospray ion source (3000 V, 35 psig, 350 °C) and mass range of 150-1000 m/z. $MH^+$ peaks were detected with the following m/z values: cGAMP 675, cdiG 691, cdiA 659, pGpA/pApG 693, pGpG 709, GMP 364, AMP 348.

**Activity Assay for GacB using Radiolabeled NTPs**

$EC_{50}$ measurements were performed using 6xHis-MBP GacB R292A, a mutant that was predicted to be an I-site mutant. This mutant was chosen to remove possible feedback inhibition during the reaction, but since then has been shown to retain cyclic di-GMP binding. GacB R292A (50 nM) was incubated in a solution of 50 mM Tris-HCl pH 7.5, 10 mM $MgCl_2$, 100 mM NaCl, and 5 mM dithiothreitol with 50 μM GTP, ~10 μCi of [$\alpha$-$^{32}$P]-GTP, and 0-5 μM of cAMP. Reaction mixtures were incubated at room temperature for 90 min., then treated with ~10 U of Calf Intestinal Alkaline Phosphatase (NEB) for 20 min at RT to degrade unreacted NTPs. An aliquot of the reaction mixture (0.5 μL) was spotted on a PEI-cellulose F thin-layer chromatography plate (Millipore), allowed to dry, then run on the plate using 1.5 M $KH_2PO_4$, pH 3.8 following the protocol developed by Kranzusch *et al* (6). Plates were dried, exposed to a Phosphor-image screen (GE Healthcare), and analyzed using a Typhoon scanner (GE Healthcare). Signal intensity was measured by ImageQuant software.

**Thermal Shift Assay**

Thermal shift assays (7) were used to assess binding between 6xHis-MBP GacB$^{1-180}$ WT GAF domain and candidate activator ligands, as well as between 6xHis-MBP GacB WT (full length) and cyclic dinucleotides. Samples were prepared at 4 °C prior to the thermal denaturation procedure. Assays were performed in a 96-well PCR plate using the CFX96 Real-Time Thermal Cycler (Bio-Rad). A reaction master mixture was prepared by mixing protein (1 μM final) and Sypro Orange (1:2500 dilution of 5000x stock, Invitrogen) in buffer containing 50 mM Tris-HCl pH 7.5, 100 mM NaCl, 10 mM $MgCl_2$, and 5 mM DTT. 49 μL aliquots were distributed to individual wells where 1 μL of ligand was added and plates were sealed. Ligands for activator screening (cAMP, adenosine, AMP, ADP, ATP, GTP, or none) were used at a final concentration of 1 μM whereas ligands for inhibition assays (cdiG, cGAMP, cdiA, or none) were used at 1 or 10 μM. Final concentrations of components in 50 μL reaction mixtures were 1 μM protein, 2x Sypro orange, and buffer containing 50 mM Tris-HCl pH 7.5, 100 mM NaCl, 10 mM $MgCl_2$, and 5 mM DTT. Reaction mixtures were simultaneously denatured with a gradient of 1 °C/min from 4-95 °C. Fluorescence was recorded using the CFX96 FRET channel, which measures broad emission signal in the absence of filters. $\Delta T_m$ values were calculated as a difference in $T_m$ between a +ligand and -ligand control.

**Bioinformatic Search for cGAMP Phosphodiesterases**

A tBLASTn search of the NCBI database nucleotide collection was performed for homologs of the PmxA HD-GYP domain (amino acids 367-562). Search parameters excluded Myxococcales and environmental samples. 14,326 of a maximum threshold of 20,000 unique sequences were returned. DNA sequences were translated using EMBOSS Transeq and multiple sequence alignments were created with MUSCLE. A Python-based program using Biopython (8) was developed to analyze alignment data of HD-GYP domains and identify sequences with specific key residues. 11,504 sequences were found to have an aligned RxxX motif. These sequences were sorted based on the identity of the fourth residue in the motif and ranked by total abundance. Sequences with Gln(Q) or Asn(N) at the fourth residue were selected and hand-curated based on conservation of active site residues, species, and lack of transmembrane regions for ease of expression and screening *in vitro*.

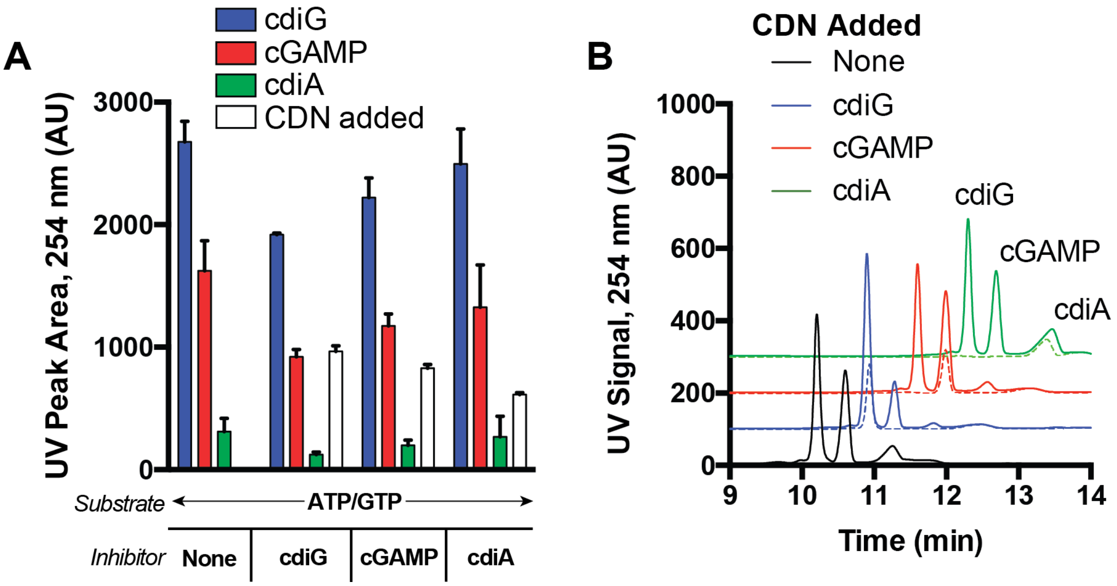


**Fig. S1. GacB is inhibited in situ by cyclic di-GMP**

(A) cdiG produced from synthase activity subtly inhibits GacB. GacB wt (0.5 µM) and CDN inhibitor candidates (10 µM) were combined with both ATP and GTP substrate (1 mM each) and reacted over 6 h. Products were measured by LC-MS and are reported as the UV peak area (254 nm). Added inhibitor signal is subtracted from the overall signal (n=2).

(B) GacB inhibition traces show product distribution in the presence of inhibitor candidates. UV traces (254 nm) of GacB inhibition in the presence of substrates and inhibitor (solid line) are overlaid with standards (dashed line) containing only 10 µM CDN (n=2).

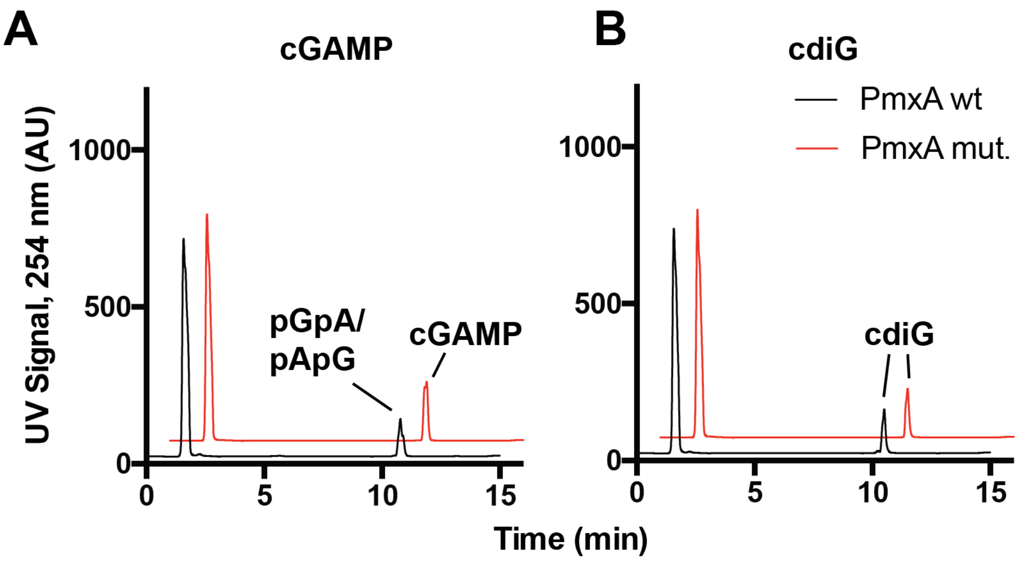


**Fig. S2. A PmxA HD-GYP mutant inactivates phosphodiesterase activity**

The conserved PmxA HD-GYP motif was mutated to AA-GYP to prohibit catalysis. 10 µM PmxA wt or mut. was incubated with 50 µM cGAMP (A) or cdiG (B) for 4 h at 30 °C. Reactions were analyzed by LC-MS. Representative LC traces for each condition are depicted, with CDN or product peaks labeled.

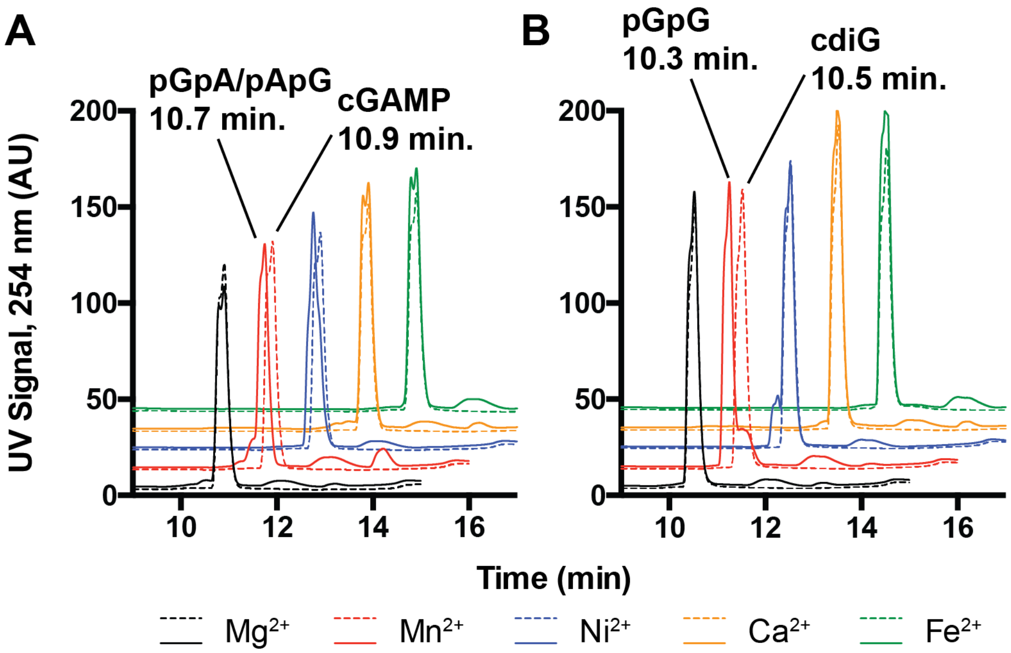


**Fig. S3. Bd is a cGAMP phosphodiesterase in the presence of $Mn^{2+}$ and $Ni^{2+}$**

10 µM Bd was incubated with 50 µM cGAMP (A) or cdiG (B) for 4 h at 30 °C. Reactions were analyzed by LC-MS and representative LC traces are shown. Five divalent cations added at 1 mM ($Mg^{2+}$, $Mn^{2+}$, $Ni^{2+}$, $Ca^{2+}$, $Fe^{2+}$) were tested for their influence on Bd activity. Reactions with Bd, CDN, and metal added (solid) are overlaid with a no enzyme control (dashed) for conditions with the same metal. Only $Mn^{2+}$ and $Ni^{2+}$ conditions yield degradation product.

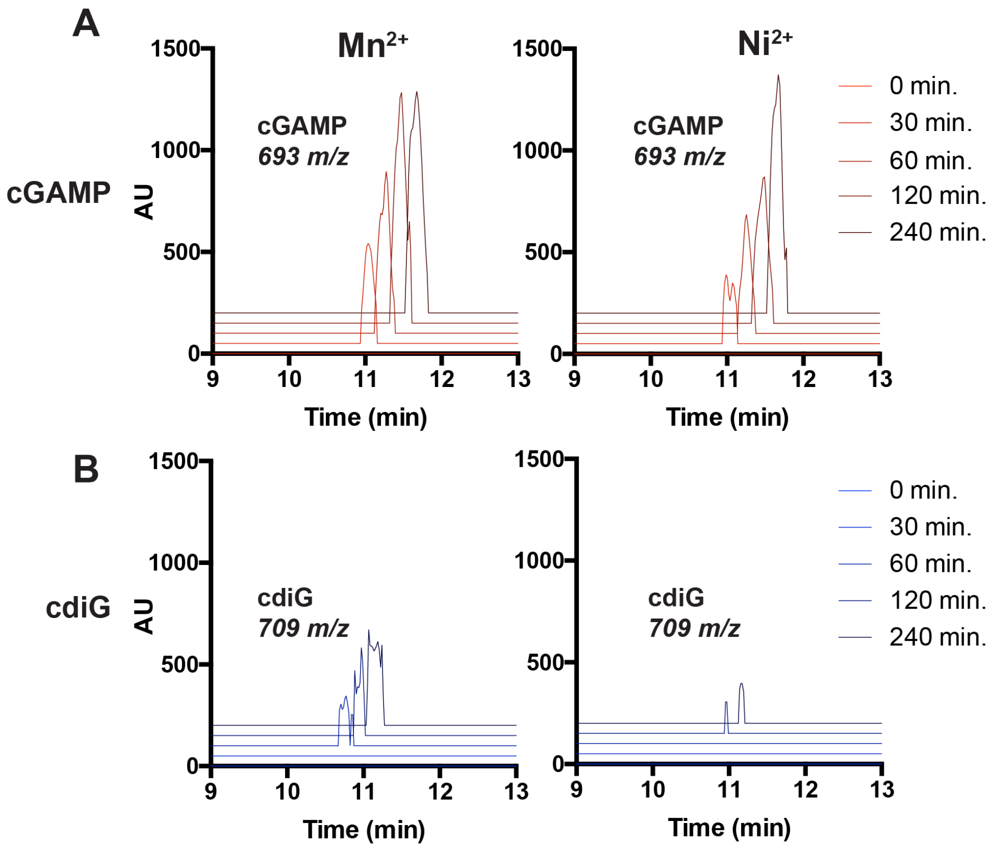


**Fig. S4. Bd is a selective cGAMP phosphodiesterase**

10 μM Bd was incubated with 50 μM cGAMP (A) or cdiG (B) for 4 h at 30 °C under conditions with $Mn^{2+}$ (left) or $Ni^{2+}$ (right), with time points measured over the interval. Results are depicted as ion extracted traces for cGAMP degradation product pApG/pGpA (red, 693 m/z) or cdiG degradation product pGpG (blue, 709 m/z). Bd with $Mn^{2+}$ displayed robust activity against cGAMP and cdiG, with a majority of either CDN degraded to the linear product within 30 minutes. Bd with $Ni^{2+}$ exhibited overall less activity, but a clear preference for cGAMP. Within 4 h, a majority of cGAMP was degraded to linear product in the presence of $Ni^{2+}$, while cdiG remained primarily as starting material.

### Table S1. Primers used in this study

| Primer Number | Sequence (5' -> 3') | Purpose |
|---|---|---|
| 1 | GAGAGGATCCATGAATCCCGCGGACCTC | Fwd Primer GacB wt, BamHI |
| 2 | GAGAGCGGCCGCCTACCGCTCCGGGAACTCGTG GGAGCCTG | Rev Primer GacB wt, NotI |
| 3 | GAGAGCGGCCGCTCACACCCGCCGGAAGTTG | Rev Primer GacB GAF domain, NotI |
| 4 | GAGAGGATCCATGTCGAAGGACGCGTACA | Fwd Primer PmxA HD-GYP domain, BamHI |
| 5 | GAGAGCGGCCGCTCAGGAGGCGAGCTTC | Rev Primer PmxA HD-GYP domain, NotI |
| 6 | CGGAAGGCCATGCCG | Fwd Primer PmxA Q518R mut. |
| 7 | GTACGGGCGGGTGG | Rev Primer PmxA Q518R mut. |
| 8 | GAGACATATGATGCGCCTGTTCAAAGC | Fwd Primer PmxA full length, NdeI |
| 9 | GAGAGGTACCTCAGGAGGCGAGCTTC | Rev Primer PmxA full length, KpnI |
| 10 | GCCATCGGGAAGATT | Fwd Primer PmxA H424A D425A mutant |
| 11 | TGCGAGGATGCCGC | Rev Primer PmxA H424A D425A mutant |
| 12 | GAGACATATGATGGGATGTCCTCCTGAC | Fwd Primer Mt, NdeI |
| 13 | GAGAAGATCTCATGGTGGCAACGCC | Fwd Primer Pp, BglII |
| 14 | GAGACATATGATGATTATTCCAGCTGATGATG | Fwd Primer Da, NdeI |
| 15 | GAGACATATGATGTATGAACTGCCGGAAAC | Fwd Primer Gm, NdeI |
| 16 | GAGACATATGATGTTTACTCAATCCCACCA | Fwd Primer Vt, NdeI |
| 17 | GAGACATATGATGGACAGCACCACGTATT | Fwd Primer Bd, NdeI |
| 18 | GAGACTCGAGTTAAGCTAAAAGGCACGATG | Rev Primer Mt, XhoI |
| 19 | GAGACTCGAGTTAGTAGTTAGGCGAAAAAGGG | Rev Primer Pp, XhoI |
| 20 | GAGACTCGAGTTAATCTTCGGAAGCACGT | Rev Primer Da, XhoI |
| 21 | GAGAGGTACCTCATACTTCCACGAAGACG | Rev Primer Gm, KpnI |
| 22 | GAGACTCGAGTTAATGACGGAATAAACGTTC | Rev Primer Vt, XhoI |
| 23 | GAGACTCGAGTTATGCTACTTTAATCTTGAACAGAA C | Rev Primer Bd, XhoI |
| 24 | GAGAGGATCCATGGACAGCACCACGTATT | Fwd Primer Bd, BamHI |
| 25 | GAGAGCGGCCGCTTATGCTACTTTAATCTTGAACA GAAC | Rev Primer Bd, NotI |
| 26 | CTTTAAGAAGGAGATATACCTATGCACAACCCTCA TGAGAGCAAGACCGACCTGGG | Fwd Primer WspR, Gibson into pETDuet-1 |
| 27 | CTTTCTGTTCGACTTAAGCATCAGCCCGCCGGGGC CGGCG | Rev Primer WspR, Gibson into pETDuet-1 |
| 28 | CTTTAAGAAGGAGATATACCTATGGAACGGATTCT CGTTGTC | Fwd Primer *Gs*GacA, Gibson into pETDuet-1 |
| 29 | CTTTCTGTTCGACTTAAGCATCAACGGATTGCCGT TGC | Rev Primer *Gs*GacA, Gibson into pETDuet-1 |
| 30 | CACGGCCGCATAATCGAAATCGATCCCGCGAAATT AATAC | Fwd Primer Dp/Gm biosensor, Gibson into pETDuet-1 |
| 31 | GGTTTCTTTACCAGACTCGACAAAAAACCCCTCAA GACCC | Rev Primer Dp/Gm biosensor, Gibson into pETDuet-1 |
| 32 | GGGTCTTGAGGGGTTTTTTGTCGAGTCTGGTAAAG AAACC | Fwd Primer pETDuet-1 backbone, Gibson |
| 33 | TCTCATGAGGGTTGTGCATAGGTATATCTCCTTCTT AAAGTTAAACA | Rev Primer pETDuet-1 backbone, Gibson |
| 34 | TGCTTAAGTCGAACAGAAAGTAATCGTATTGTACA CGGCCGCATAATCGAAAT | Fwd Primer Extend biosensor and add overhang with synthase Rev Primer |

**Table S2. Protein sequences used in this study**

| Gene | Sequence |
|---|---|
| GacB wt | MNPADLLSAMKRTVEQLAAFNEMAKALTSTLELREVLALVMQKVSSLLLPRNWSLILQDERTGKLYFEIAVGDGADVLKGLQLNPGEGIAGAVFTSGAARLVHDVGGDPSFSPRFDEASAFHTRSILAVPLLARGRVLGIIELVNGPMDPPFTNEDLTILTAIADYAAIAIENARNFRRVQELTITDEHTGCYNARHLRALLDQEVKRSERFSHPLSLVFLDLDHFKSINDTHGHLVGSATLKEVGDLLMTLGRQNLDAVFRYGGDEFAMLLVETDPEGAAVIGQRVCEAFRGRGFLLEQGLDVRLTASVGVATYPDHASSALDLIRAADFAMYAAKARGRDALCIAEPIAPNGGTGSHEFPER* |
| GacB GAF domain | MNPADLLSAMKRTVEQLAAFNEMAKALTSTLELREVLALVMQKVSSLLLPRNWSLILQDERTGKLYFEIAVGDGADVLKGLQLNPGEGIAGAVFTSGAARLVHDVGGDPSFSPRFDEASAFHTRSILAVPLLARGRVLGIIELVNGPMDPPFTNEDLTILTAIADYAAIAIENARNFRRV* |
| PmxA HD-GYP domain | MSKDAYTRGHSQRVGDVSVQIGREMKLTERELRQLQYGGILHDIGKIGIVESILCKQTRLTDQEMDIMREHPAIGDAIIGPVSFLGAVRACVRHHHERWDGTGYPDKLKGEDIPLLARIVACADTFDACTSTRPYQKAMPLEKAMEILDNLSGAQLDPKVVQALKQVLAKQGVRLEGHRLPVKLAS* |
| PmxA HD-GYP domain Q518R | MSKDAYTRGHSQRVGDVSVQIGREMKLTERELRQLQYGGILHDIGKIGIVESILCKQTRLTDQEMDIMREHPAIGDAIIGPVSFLGAVRACVRHHHERWDGTGYPDKLKGEDIPLLARIVACADTFDACTSTRPYRKAMPLEKAMEILDNLSGAQLDPKVVQALKQVLAKQGVRLEGHRLPVKLAS* |
| PmxA HD-GYP domain H424A D425A | MSKDAYTRGHSQRVGDVSVQIGREMKLTERELRQLQYGGILAAIGKIGIVESILCKQTRLTDQEMDIMREHPAIGDAIIGPVSFLGAVRACVRHHHERWDGTGYPDKLKGEDIPLLARIVACADTFDACTSTRPYQKAMPLEKAMEILDNLSGAQLDPKVVQALKQVLAKQGVRLEGHRLPVKLAS* |
| PmxA wt (full length) | MRLFKAILLLMLVVGVVPTLMVGWLSVSHTRELLVRDAQELAQERVKQLRLKAEIFLGEPTDAVLGLARVPGFFGLPLEAQQTHLASVLNQRRDVLALTVFDANRQRLPGLQAFSKHDVPPTALAGHEERARALLENIEGVRYADVVKDPQGSEPVLTLAFPLGEPVRGYIAADLSLADLRKMLEQERVGSTGFAYLADRHGHLVAGGGGVAGLGEDVAQRGPVAHLLKQLVGKSDMELFHVGNFGEGRDAVVAAYTVLPETGWAIVSEQPVEHAYRQVDTMEQRILLGLGAAILVAVVLAALFSRNLTRPLKGFISGALELAHGKFGVEVDIKQKNELGELAQTFNYMSKQLLAYDMENRGLYESLEKGYLETIVALANSIDSKDAYTRGHSQRVGDVSVQIGREMKLTERELRQLQYGGILHDIGKIGIVESILCKQTRLTDQEMDIMREHPAIGDAIIGPVSFLGAVRACVRHHHERWDGTGYPDKLKGEDIPLLARIVACADTFDACTSTRPYQKAMPLEKAMEILDNLSGAQLDPKVVQALKQVLAKQGVRLEGHRLPVKLAS* |
| PmxA H424A D425A (full length) | MRLFKAILLLMLVVGVVPTLMVGWLSVSHTRELLVRDAQELAQERVKQLRLKAEIFLGEPTDAVLGLARVPGFFGLPLEAQQTHLASVLNQRRDVLALTVFDANRQRLPGLQAFSKHDVPPTALAGHEERARALLENIEGVRYADVVKDPQGSEPVLTLAFPLGEPVRGYIAADLSLADLRKMLEQERVGSTGFAYLADRHGHLVAGGGGVAGLGEDVAQRGPVAHLLKQLVGKSDMELFHVGNFGEGRDAVVAAYTVLPETGWAIVSEQPVEHAYRQVDTMEQRILLGLGAAILVAVVLAALFSRNLTRPLKGFISGALELAHGKFGVEVDIKQKNELGELAQTFNYMSKQLLAYDMENRGLYESLEKGYLETIVALANSIDSKDAYTRGHSQRVGDVSVQIGREMKLTERELRQLQYGGILAAIGKIGIVESILCKQTRLTDQEMDIMREHPAIGDAIIGPVSFLGAVRACVRHHHERWDGTGYPDKLKGEDIPLLARIVACADTFDACTSTRPYQKAMPLEKAMEILDNLSGAQLDPKVVQALKQVLAKQGVRLEGHRLPVKLAS* |

| | |
|---|---|
| Moth_2495 (Mt) | MGCPPDRSFCELVVALSTILDIEEETKLYHAWRVALVAQELARRVIPDEATLVFYGGLLHDIGAMGLD<br>DHLVHLALQRGSRNNPEVVNHPLRGADMVAAIPGLGEKVAAMIRDHHERWNGSGYPRGIAGNHIVT<br>GAMLLGLADELDLVLRVHPGTSWARLRETLNRRVQGGFPPELLAILDKMMNGPLYAEIATNTALELK<br>MFKVILDLPPINFQVPDPMKITIDLFARIIDAKHAYTAGHSHRVAAYALNLARCLGYNDAKLRRLEIAGL<br>LHDFGKIAVPRAILDKRGRLNSEELKVVRRHPAWTIELLEGVTSLKDIARDAGLHHERYDGKGYPYGL<br>HDGEIPLGARIIAVADAFDAMTSNRPYQPTRTPEEALKILAGGAGTQFDPEVTAVASCLLA* |
| PhaeoP14_01534 (Pp) | MVATPTPYSLPGPDPGPGFAAASTGVLHDVTYPRVGGQTIQLGELLGALSHALDLTEGQPRGHCVR<br>CCWIGMQVADYLHLSRQDRSDLYFTLLLKDLGCSSNAARISTLYRTDDLSFKRNFKFVDDTTRKKLD<br>FLLQHTGQGDSGFKRLKTVTGVIARSDSIATELIQTRCDRGAKIARLMRFSERIAGGIAALDEHWNGG<br>GHPAGISRKDIPLFSRIALMAQVTDVFALQYGSGAAADELESRAGSWFDPELVPVFTRVIRTPGFYDT<br>LTSDRLETSVFSDPAARATLGVDDTYMDDISYAFSKVIDAKSPFTYGHSERVAHYTDMICETLGFELP<br>HRRWMVRAGLLHDLGKLGVSNAILDKPGRLSDQEMKQMKLHPEFGYEVLRRISVFQDIADVVVAHH<br>ERLDGKGYPYGLGAADLTVEMRILTVADIFDAMTADRPYQSALPLEKAFATMDEMQGTAIDPEIHRAL<br>KEAIAASPWPAREARPFSPNY* |
| Daro_3290 (Da) | MIIPADDATPIDTIAHALTVALGARDHYTRIHCDRVVNLSAELGQHIGLNSEELERLSLGARFHDLGKIG<br>IPDIVLRKPEPFEAHEWECMKQHVLIGEQIVLAINGGRASRIARTVRHHHEHFDGSGYPDGLTGSQIP<br>LDARIISVVDSYDAMIARRPYQNARDHQAVMDILISESGTKHDPDLVHAFGAIIEKSASRASED* |
| Gmet_3476 (Gm) | MYELPETAPVRVLFVDDEENMLKSLSRVFMDEEFEVITAPSAREGLRILMETEDVGVIVSDQRMPEIT<br>GVQFLAQVREMFPETLRIMLTGYADISATVDAINRGGAYRYITKPWNDEELVQIIRDAVNQYRLVQEN<br>RRLTEIVNKQNEELKEWNENLKVRVLEQTTEIRKKHENLKELNERLRSNFKDTVTAFSRLIELGGKKR<br>HAETVATLAMNIAQAMGVSAGEVETVHNAALLHDIGELGIADAILKKDLSEMTPGELREYMLHAVRG<br>QTAIDAIEELTPAGVLIRHHHEHYDGSGYPDGLRGEAIPLGSRIIAIADFLDRAIQKLHGYTAIDYGLRS<br>VAAELGKKLDPSLYESLRKFAKHVYYVKETDTRKLGEVEVGIAELFVGQVLSRDIYSGTGLLLLNRGV<br>RLDVVKIEAIRRYYRLDPPPHGVFVEV* |
| Bd2325 (Bd) | MDSTTYFRIRLSTIRPDKVTSFDIYILVDGKHILYLRAGDKLASGKIKTLHGRDTGDSFFVRIEDKQTYR<br>DWVKEEMNSDLLNPFDKAKILRESSVALMEDLFENPDVNKALDESRPIITDFIDLMENAPEAMGFMIS<br>LSGHDFYTYNHSLDVSIYSLGLGKALGYDAKTLEELGVGALFHDIGKRSVSLDILCKKGGLTDAEWEQ<br>MKMHPQYGLVIMNNHPNISDAIKAACFEHHESWAGNGYPQQLVAEEIHPFARIVAITDTYDAMTTQR<br>SYNVPMTPMDAVAMMKEKLAGRYDPDMLKAMYSVLFKIKVA* |
